# A prevalent and culturable microbiota links ecological balance to clinical stability of the human lung after transplantation

**DOI:** 10.1101/2020.05.21.106211

**Authors:** Sudip Das, Eric Bernasconi, Angela Koutsokera, Daniel-Adrien Wurlod, Vishwachi Tripathi, Germán Bonilla-Rosso, John-David Aubert, Marie-France Derkenne, Louis Mercier, Céline Pattaroni, Alexis Rapin, Christophe von Garnier, Benjamin J. Marsland, Philipp Engel, Laurent P. Nicod

## Abstract

There is accumulating evidence that the lower airway microbiota impacts lung health. However, the link between microbial community composition and lung homeostasis remains elusive. We combined amplicon sequencing and culturomics to characterize the viable bacterial community in 234 longitudinal bronchoalveolar lavage samples from 64 lung transplant recipients and established links to viral loads, host gene expression, lung function, and transplant health. We find that the lung microbiota post-transplant can be categorized into four distinct compositional states, ‘pneumotypes’. The predominant ‘balanced’ pneumotype was characterized by a diverse bacterial community with moderate viral loads, and host gene expression profiles suggesting immune tolerance. The other three pneumotypes were characterized by being either microbiota-depleted, or dominated by potential pathogens, and were linked to increased immune activity, lower respiratory function, and increased risks of infection and rejection. Collectively, our findings establish a link between the lung microbial ecosytem, human lung function, and clinical stability post-transplant.

**Graphical abstract:** 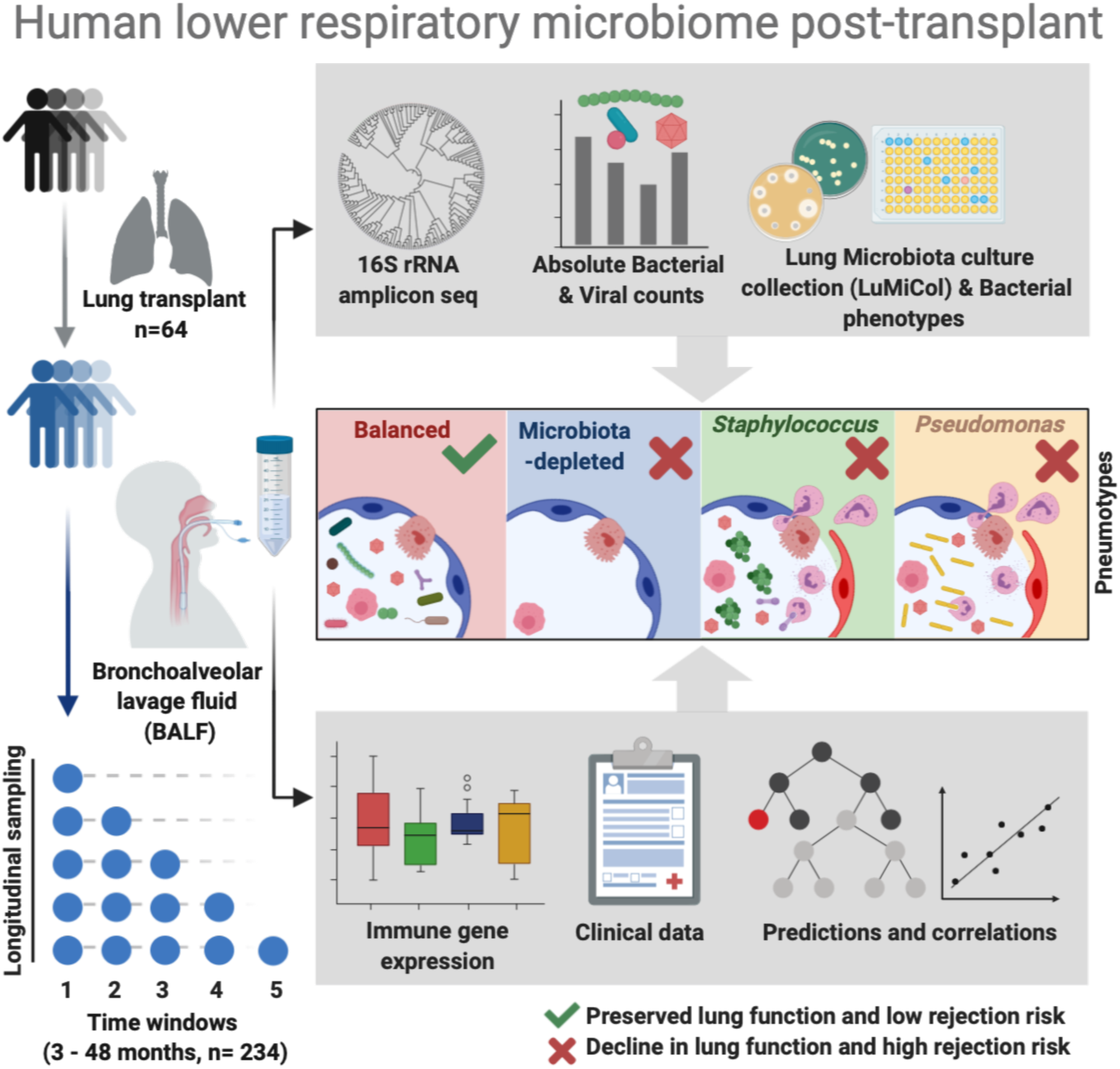

## Introduction

Recent studies have shown that diverse bacterial communities are present in the lower respiratory tract of healthy humans (Charlson et al., 2011; Dickson et al., 2015, 2017; Pattaroni et al., 2018; Segal et al., 2013; Venkataraman et al., 2015). These communities are predominated by the same phyla as the oral and gastrointestinal microbiota (*Bacteroidetes*, *Firmicute*s, Actinoabacteria, Proteobacteria). However, their phylogenetic composition, total bacterial load, and temporal-spatial dynamics are distinct owing to the characteristic physicochemical, anatomical, and immunological conditions of the lung, which makes this organ a distinct microbial habitat with specific host-microbe interactions (Dickson et al., 2015; Lloyd and Marsland, 2017).

Several independent studies have shown that supraglottic taxa (i.e bacteria found in the human oropharyngeal area) such as *Streptococcus, Prevotella*, and *Veillonella* are major constituents of the healthy lower respiratory tract microbiota. These bacteria have been proposed to contribute to the immunological development and homeostasis of the human lung, as their presence correlates with an increased pro-inflammatory response during postnatal immune maturation as well and lung function in adulthood (Pattaroni et al., 2018; Segal et al., 2016). Shifts in microbial community composition, characterized by decreased bacterial diversity and collectively referred to as “dysbiosis” (Dickson and Huffnagle, 2015; Marsland and Gollwitzer, 2014), have been associated with various respiratory diseases such as Chronic Obstructive Pulmonary Disease (COPD), Idiopathic Pumonary Fibrosis (IPF) and asthma. Together, these findings suggest that the lower respiratory microbiota is linked to the health state of the human lung and hence may play important roles for maintaining lung homeostasis.

Formidable challenges are associated with studying the lower respiratory tract microbiota. Firstly, the sampling of the human lung, which is best achieved by collecting bronchoalveolar lavage fluid (BALF) during bronchoscopy (Carney et al., 2020), is an invasive procedure which implies that it is rarely performed in healthy individuals. Consequently, large datasets from healthy individuals - including longitudinal studies that would inform about the dynamics of the human lung microbiota - are scarce. Secondly, the relatively low bacterial biomass in the human lung increases the risk of describing contaminants as being part of the respiratory tract microbiota. This can skew diversity measures of the lower respiratory tract microbiota, in particular when solely relying on relative abundance data (Segal et al., 2013). Thirdly, only few studies haves attempted to isolate viable bacteria from the human lung (Cummings et al., 2020; Venkataraman et al., 2015; Whelan et al., 2020), and little is known about their physiology and growth characteristics. Therefore, our current understanding of the ecological properties of different lung microbiota members and how these are linked to the environmental conditions in the lung ecosystem (such as immune state) remains limited.

Studying the microbiota in the context of lung transplantation can provide important insights about the crosstalk between the respiratory tract microbiota and the host (Mouraux et al., 2017). Lung transplant recipients undergo post-transplant follow-up, in which BALF is collected to monitor the health state of the transplanted organ. This offers unique opportunities for longitudinal studies on the lung microbiota composition and allows establishing links to the host’s immune state and to clinical metadata. Due to different types of clinical complications such as infection (Nosotti et al., 2018), acute cellular or humoral rejection (Martinu et al., 2011) and Chronic Lung Allograft Dysfunction (CLAD) (Koutsokera et al., 2017), the transplanted lung also offers the opportunity to study the respiratory microbiota (Borewicz et al., 2013; Charlson et al., 2012; Gregson et al., 2013; Willner et al., 2013) under a wide variety of ecological conditions. A better understanding of the dynamics of the lung ecosystem in this context can ultimately help limit the burden of morbidity and mortality associated with post-transplant complications and promote graft survival.

Recent studies on lung transplants have provided insights about the distribution of the microbiota along the conducting and respiratory airways (Beaume et al., 2016), or the adaptation of opportunistic pathogens to the lung environment (Beaume et al., 2017). Moreover, there is accumulating evidence that the immune state of the transplanted lung correlates with changes in the composition of the lung microbiota (Bernasconi et al., 2016; Charlson et al., 2012). High abundance of opportunistic pathogens such as members of the *Staphylococcus* and *Pseudomonas* genera have been linked to pro-inflammatory responses in the transplanted lung (Bernasconi et al., 2016; Erb-Downward et al., 2011), and also found in respiratory diseases such as COPD and asthma (Hilty et al., 2010; Mika et al., 2018). These bacteria activate macrophages and induce a strong inflammatory response after transplantation, reflected by high levels of tumor necrosis factor-α (TNFα) and cyclooxygenase-2 (COX2)(Bernasconi et al., 2016). This is in contrast to taxa such as *Streptococcus*, whose abundance has been linked to low inflammation and tissue repair and remodeling (Bernasconi et al., 2016). Sustained inflammatory reactions and uncontrolled tissue remodeling can eventually lead to irreversible decline in lung function (Hardison et al., 2009; Todd et al., 2020). These previous data collectively suggest that the lung microbiota post-transplant can constitute different compositional states that may be linked to allograft function. However, quantitative analysis of these microbiota profiles are currently lacking, including the phylogenetic and physiological characterization of viable community members, and the links to the lung ecological environment and the clinical outcome post-transplant.

In this study, we characterized the airway microbiota in 234 longitudinal BALF samples from 64 lung transplant recipients. We combined culture-independent and -dependent analysis to identify the most prevalent lung bacteria post-transplant and to establish a strain collection of primary lung bacterial isolates. We linked the identified compositional changes in lung microbiota to host gene expression profiles, anellovirus loads and patient metadata to understand the importance of the ecological environment of the transplanted lung on clinical outcomes. Our findings show that BALF samples can be classified into four distinct compositional states (i.e. pneumotypes) similar to the enterotypes identified in the human gut (Arumugam et al., 2011). These pneumotypes are distinguished by different community characteristics and distinct physiological properties of their predominant members. We show that pneumotypes are differentially associated with anellovirus loads, respiratory function, and both local and peripheral host immune responses, including those linked to allograft rejection. Taken together, our findings not only illustrate the strong links between lung health and local microbiota composition, but pinpoint underlying community characteristics and lung environmental conditions as well as provide a large resource of cultured isolates for future experimental approaches

## Results

### Combined culture-dependent and -independent approach identifies the prevalent and viable bacterial community members of the human lung post-transplant

To characterize the bacterial community composition of the lung microbiota post-transplant, we performed 16S rRNA gene amplicon sequencing of 234 longitudinal BALF samples from 64 lung transplant recipients collected over a 49-month period (**Figure 1A, Table S1**). A total of 7,164 operational taxonomic units (OTUs) were identified, excluding OTUs contributing to reads in 11 negative control samples (See **Methods, Figure S1A, Dataset S1, S2**. In accordance with previous studies on BALF samples from healthy non-transplant individuals (Erb-Downward et al., 2011; Pattaroni et al., 2018; Segal et al., 2013; Venkataraman et al., 2015), we found that *Bacteroidetes* and *Firmicutes* followed by *Proteobacteria* and *Actinobacteria* are the most abundant phyla in the post-transplant lung (**Figure 1B**). Prevalence analysis across all BALF samples showed that the community composition is highly variable with only 22 OTUs shared by ≥50% of the samples (**Figure S1B, Dataset S3)**. However, these 22 OTUs constituted 42 % of the total number of normalized reads, indicating that they are predominant members of the post-transplant lung microbiota (**Figure 1C, Figure S1C, Table S2, Dataset S3**). They belonged to the genera *Prevotella7*, *Streptococcus*, *Veillonella, Neisseria, Alloprevotella, Pseudomonas, Gemella, Granulicatella, Campylobacter, Porphyromonas and Rothia*, the majority of which are also prevailing community members in the healthy human lung (Dickson et al., 2015, 2017; Erb-Downward et al., 2011; Segal et al., 2013), suggesting a considerable overlap in the overall composition of the lung microbiota between the healthy and the transplanted lung.

**Figure 1.**
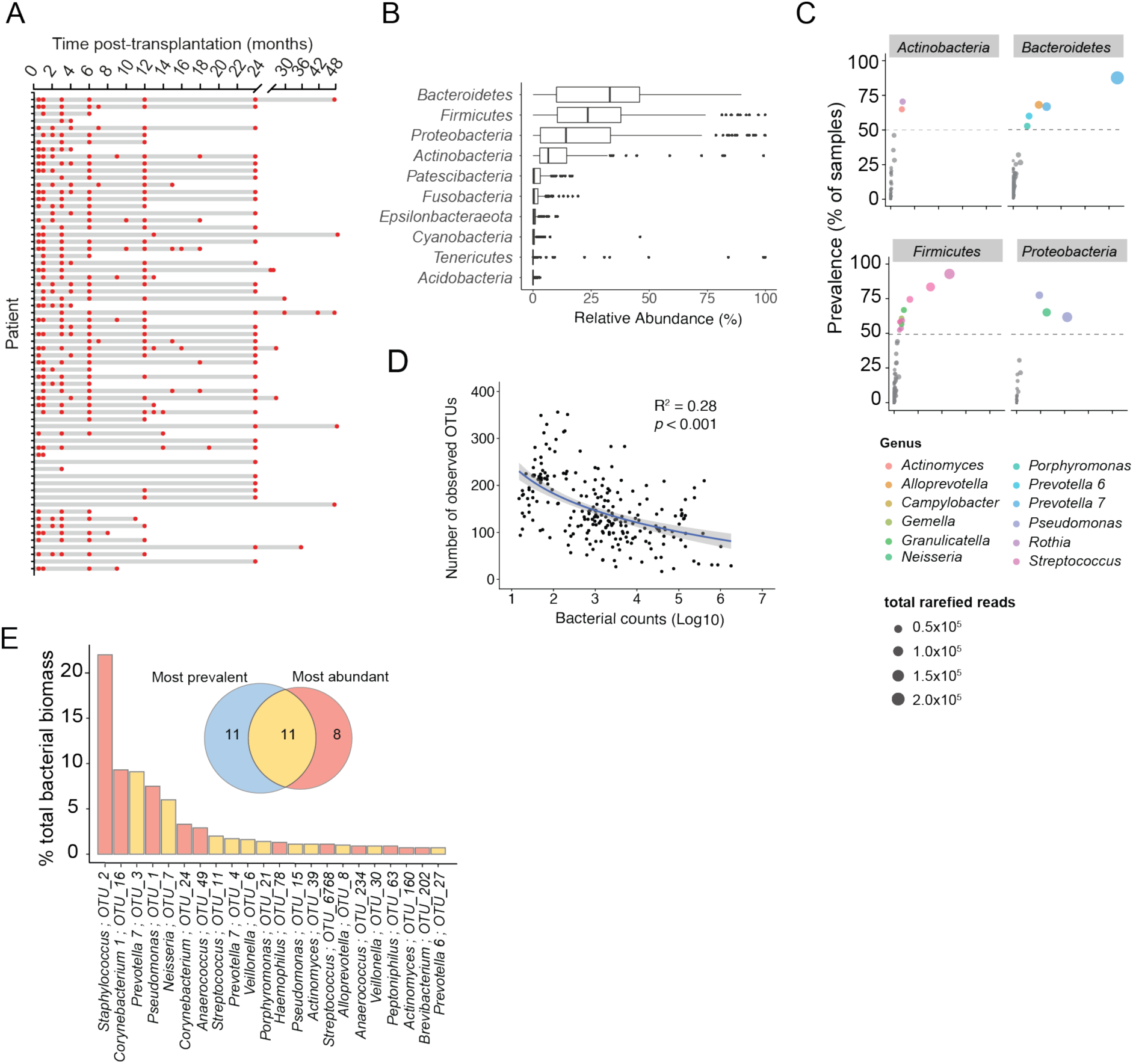
Combining BALF amplicon sequencing and culturomics to deduce the microbial ecology of deep lung microbiota. **(A)** Schematic of longitudinally obtained Bronchoalveolar lavage fluid (BALF) from lung transplant recipients over time (months post-transplant, See Methods). **(B)** Median relative abundances (%) of most abundant phyla across BALF samples are plotted as box plots. **(C)** Prevalence (≥50% of samples - grey dashed line) vs contribution to total normalized reads (dot size) across samples for most abundant phyla and genera (colored dots). **(D)** Correlation between number of OTUs and bacterial counts detected per BALF sample **(E)** Bacterial taxa (genera; OTU IDs) contributing ≥ 75% of total bacterial counts (%) plotted as bar chart. Venn diagram inset shows overlap (yellow) between the most prevalent (≥50% incidence, light blue) and the most abundant (≥75% total count, red) taxa in the transplanted lung. Bar colors denote the categories to which represented taxa belong. Errors bars indicate median ± interquartile range. Bacterial count / sample was obtained by quantifying 16S rRNA gene copies with qPCR. R^2^ was given by linear regression, *p*< 0.05. Contribution of OTUs to total bacterial counts across all samples was obtained as the Bacterial counts / sample x n i.e. 234.

Differences in bacterial loads between samples can skew community analyses when based on relative abundance profiling alone. Therefore, we used qPCR to determine the total copies of the 16S rRNA gene as an estimate for bacterial counts, and normalized the abundances of each OTU across the 234 samples (absolute abundance). We found that the bacterial counts vastly differed between samples, ranging between 10^1^ and 10^6^ gene copies per ml of BALF (**Figure S1D**). The number of observed OTUs increased with decreasing counts (**Figure 1D**) suggesting that a large fraction of the OTUs were detected in samples of low bacterial biomass and hence represent either transient or extremely low-abundant community members, or sequencing artefacts and contaminations. In turn, 19 of the 7,164 OTUs constituted >75% of the total absolute abundances detected across the 234 BALF samples (**Figure 1E**). This included 11 of the 22 most prevalent OTUs (see above) plus eight OTUs that were detected in only a few samples but at very high abundance (*Staphylococcus ; OTU_2, Corynebacterium 1; OTU_16 and OTU_24, Anaerococcus; OTU_49* and *OTU_234, Haemophilus; OTU_78, Streptococcus; OTU_6768, Peptoniphilus; OTU_63,* **Table S2**). It is important to differentiate these opportunistic colonizers from other community members with low incidence, as they reached very high bacterial counts in some samples with potential implications for lung health.

To demonstrate the viability of prevalent lung microbiota members and to establish a reference catalogue of bacterial isolates from the human lung for experimental studies, we complemented the amplicon sequencing with a culturomics approach (**Figure S2**). We cultivated 21 random BALF samples from 18 individuals, on 15 different semi-solid media (both general and selective) in combination with 3 oxygen concentrations; aerobic, 5% CO_2_, and anaerobic (**Dataset S4, Methods**), representing 26 different conditions. This resulted in a total of 300 bacterial isolates, representing 5 phyla, 7 classes, 13 orders, and 17 families from which we built an open-access biobank of bacterial isolates, called the **Lu**ng **Mi**crobiota culture **Col**lection of bacterial isolates (LuMiCol, **Dataset S5**)

To examine the extent of overlap between bacteria in LuMiCol and the diversity obtained by amplicon sequencing, we included 16S rRNA gene sequences from 215 isolates that passed our quality filter into the community analysis, which allowed for the retrieval of OTU-isolate matching pairs (Methods). We cultured fresh BALF immediately upon extraction (within 2 hours), as we observed loss in bacterial diversity upon cultivating frozen samples. We found that 213 isolates matched to 47 OTUs (**Figure 2A-C, Dataset S6**), including 17 of the most prevalent and abundant bacteria (Figure 1E, **Table S2**). As expected, bacteria with high abundance in the amplicon sequencing-based community analysis were isolated more frequently, with *Firmicutes* revealing the highest isolate diversity (**Figure 2A-C, Datasets S5, S6**) and being recovered under the most diverse culturing conditions.

**Figure 2.**
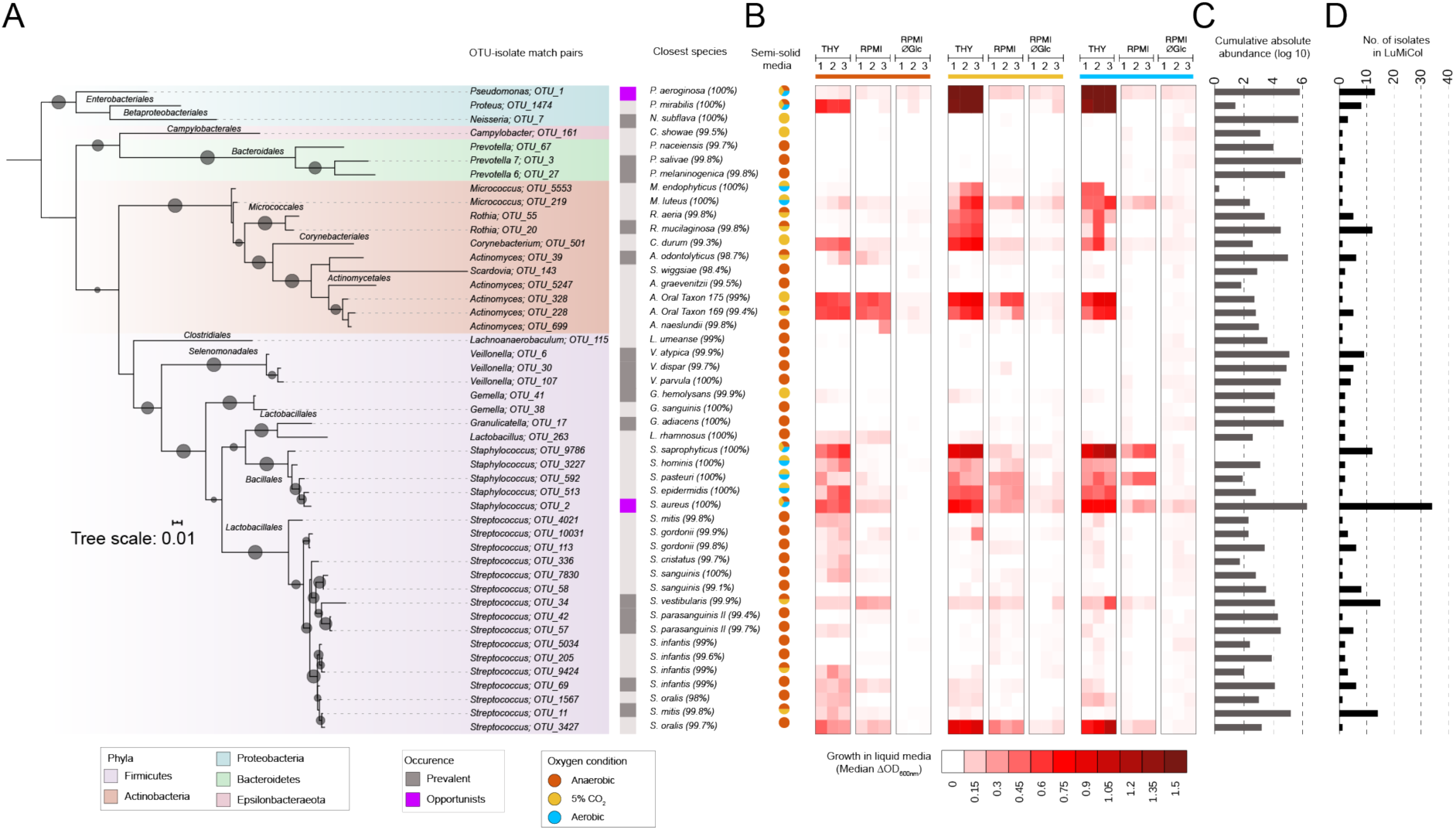
A lung microbiota culture collection (LuMiCol) reveals extended diversity and phenotypic characteristics of the lower airway bacterial community. **(A)** Phylogenetic tree of the 47 OTU-isolate matching pairs inferred with FastTree (See Methods). Branch boot strap support (size of dark grey circles) ≥80% is displayed. **(B)** Growth characteristics of each OTU-isolate matching pair in three different oxygen conditions (Anaerobic - light brown, microaerophilic-yellow, aerobic-light blue, n= 3). Column with pie charts shows growth on semi-solid agar. Heatmap shows median change in Optical Density (OD) at 600 nm growth in three different liquid media (THY, RPMI, RPMI without glucose) over three days. **(C, D)** Cumulative counts of each OTU-isolate matching pair across all BALF samples (grey) and the number of isolates in Lumicol (black) are plotted as bars. Taxa are labeled as genus ; OTU ID, with an indication of whether they are prevalent (grey rectangle) or opportunistic (magenta rectangle) in the lower airway community. The names of the closest hit in databases: eHOMD and SILVA are used as species descriptor.

In summary, our results from the combined culture and culture-independent approach show that the lung microbiota post-transplant is highly variable in terms of both bacterial load and community composition with many transient and low-abundant bacterial taxa. However, a few community members display relatively high prevalence and/or abundance suggesting that they represent important colonizers of the human lung.

### LuMiCol informs on the diversity and metabolic preferences of culturable human lung bacteria

We characterized the culturable community members of the lower respiratory tract contained in LuMiCol by testing a wide range of growth conditions and phenotypic properties (**Methods)**. The majority of the cultured isolates could taxonomically be assigned at the species level based on genotyping of the 16S rRNA gene V1-V5 region. However, the limited taxonomic resolution offered by this method does not allow to discriminate between closely related strains, which can include both pathogenic and non-pathogenic members. Hence for *Streptococcus,* we additionally tested for type of hemolysis (alpha, beta, or gamma) and resistance to optochin, which differentiates the pathogenic pneumococcus and the non-pathogenic viridans groups (**Figure 2A, Figure S2B, C**). This demonstrated that the 16 matched pairs of *Streptococcus* OTU-isolate pairs belong to the viridans group of Streptococci (VS)(Bowers and Jeffries, 1955). Interestingly, these isolates exhibited the highest genotypic and phenotypic diversity throughout our collection and belonged to five OTUs among the 22 most prevalent community members, with *Streptococcus mitis* (OTU_11) present in 93.6% of all samples.

BALF from healthy individuals contains amino acids, citrate, urate, fatty acids, and antioxidants such as glutathione but no detectable glucose (Evans et al., 2014), which is associated with increased bacterial load and infection (Brennan et al., 2007; Gill et al., 2016; Mallia et al., 2018). To get insights into basic bacterial metabolism, we assessed the growth of all 47 isolates matching an OTU under different oxygen concentrations. In addition to the different conditions used during isolation on semi-solid media, we also used undefined rich media (Todd-Hewitt Yeast extract) and defined low-complexity liquid media (RPMI 1640), for which we also used a glucose-free version to mimic the deep lung environment (**Methods**). Despite the presence of oxygen in the human lung, the majority of the isolates were either obligate or facultative anaerobes (**Figure 2A),** including some of the most prevalent members (*Prevotella melaninogenica (OTU_3), Streptococcus mitis*; *OTU_11, Veillonella atypica (OTU_6)* and *Granulicatella adiacens (OTU_17)*. A similar trend was also observed in liquid media under anaerobic conditions, with the exception of the genera *Prevotella*, *Veillonella* and *Granulicatella.* Most Streptococci from the human lung grew best in complex media containing glucose under anaerobic conditions, including the most prevalent bacteria in our cohort, *S. mitis* (OTU_11) (**Figure 2B**). However, noticeable exceptions were *S. vestibularis* (OTU_34), *S. oralis* (OTU_3427 and OTU_1567), *S. gordonii* (OTU_10031), which grew equally well in the presence of oxygen and in low-complexity medium (**Figure 2B**). Most *Actinobacteria* grew best on rich medium under microaerophilic conditions (5% CO_2_), with an exception of *A. odontolyticus* (OTU_39), which required anaerobic conditions. Some *Actinobacteria* grew equally well under anaerobic conditions i.e. *C. durum* (OTU_501), *Actinobacteria* sp. oral taxon (OTU_328 and OTU_228).

The two most predominant opportunistic bacteria in our lung cohort, *P. aeruginosa* (OTU_1) and *S. aureus* (OTU_2), grew best in rich media in the presence of oxygen, although the latter also grew under anaerobic conditions (**Figure 2C**). Although *in vitro*, these results indicate towards changes physicochemical conditions in the lung that may favor the growth of aerobic bacteria with potentially pathogenic properties. In summary, our insights from the bacterial culture collection provide first insights into the phenotypic properties of human lung bacteria and will serve as a basis for future experimental work.

### Identification of four compositionally distinct pneumotypes post-transplant using machine learning based on ecological metrics

To detect and characterize differences in bacterial community composition between BALF samples from transplant patients, we clustered the samples using an unsupervised machine learning algorithm based on pairwise Bray-Curtis dissimilarity (beta diversity, **See Methods**). This segregated the samples into four partitions around medoids (PAMs) at both phylum and OTU level (**Figure 3A, B, S3A, S3B**). We refer to these clusters as “pneumotypes” PAM1, PAM2, PAM3, and PAM4 (**Table 3**). PAM1 formed the largest cluster consisting of the majority of samples (n=115) followed by PAM3 (n=76), PAM2 (n=19), and PAM4 (n=24) (**Dataset S8**). Examination of various diversity measures (OTU richness, OTU diversity, Species occurrence, **Figure 3C-E**), distribution of the dominant community members (**Figure 3F**), and bacterial counts (16S rRNA gene copies, **Figure 3G**) revealed distinctive characteristics between the four pneumotypes.

**Figure 3.**
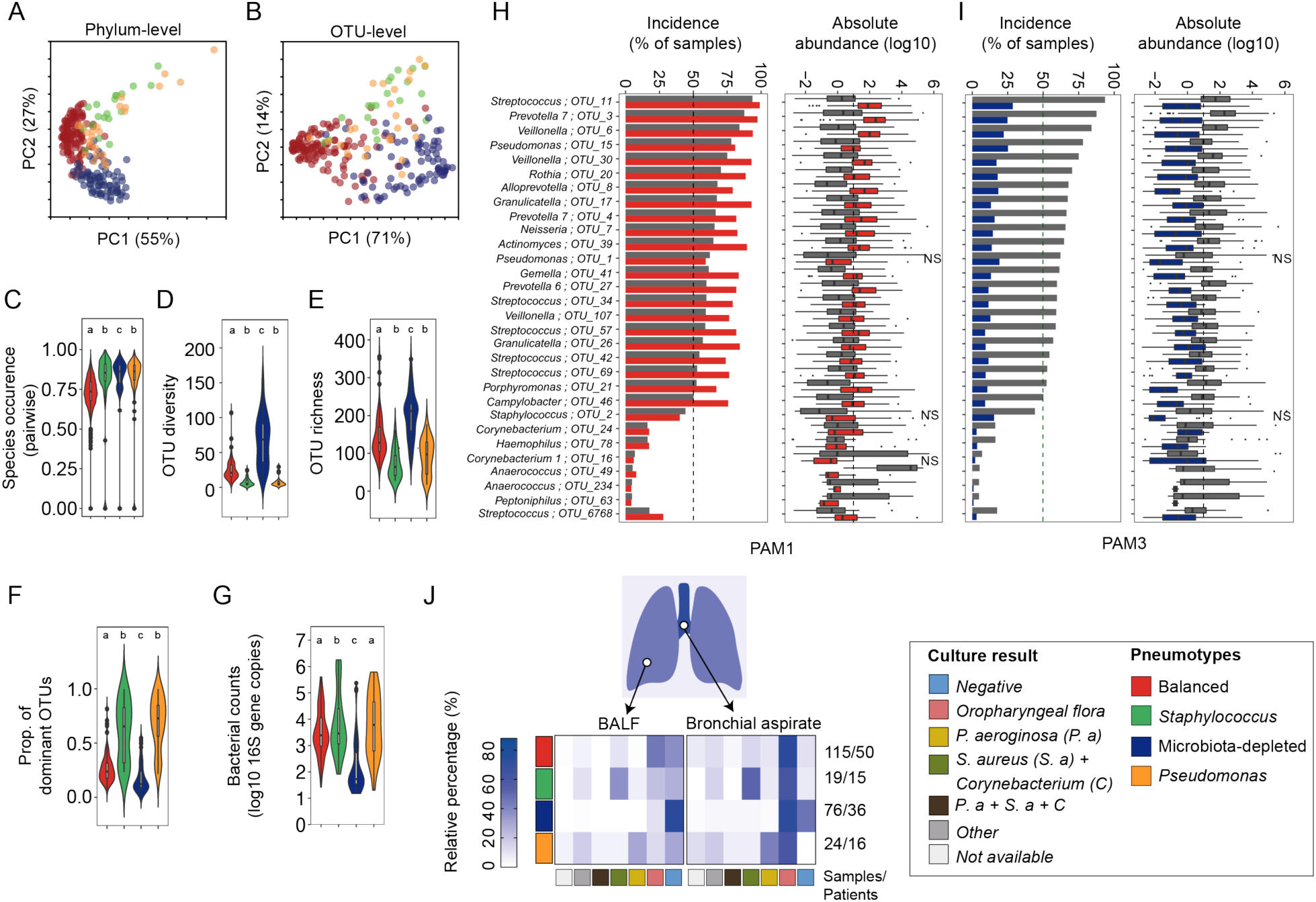
Bacterial communities of the lung post-transplant fall into four ‘pneumotypes’ with distinct community characteristics. **(A, B)** Principal component analysis shows Partition around medoids (PAMs) at phylum and OTU level respectively (See also **Figure S3)** generated by K-medoid-based unsupervised machine learning using Bray-Curtis dissimilarity (occurrence and abundance). **(C-G)** Violin plots (with inset boxplots) showing pairwise species occurrence (Sorenson’s index), OTU diversity, OTU richness, proportion of most dominant OTUs and total bacterial counts, respectively, across 4 pneumotypes (one-way ANOVA with Tukey’s post hoc test or Kruskal-Wallis test with Dunn’s post hoc test, *p*< 0.05). **(H and I)** Enrichment analysis of prevalence (See methods, ≥50% - green dotted line) and absolute abundance across all samples of the 30 most dominant taxa (i.e. OTUs) in Pneumotype_balanced_ and Pneumotype_MD_ respectively, when each was compared to the other 3 combined (grey, See also Figure S4). **(J)** Heatmap shows relative percentage of taxa (right colored panel) cultured from paired samples of Bronchial aspiration (BA) and Bronchoalveolar lavage fluid (BALF) from each pneumotype (left colored panel). Oropharyngeal flora mainly corresponds to Pneumotype_balanced_ (i.e. *Streptococcus, Prevotella*, *Veillonella*). Alpha diversity calculated using Renyi diversity with corresponding Hill numbers (See Methods). Pneumotypes are color coded: Balanced (red), *Staphylococcus* (green), Microbiota-depleted (MD, blue), and *Pseudomonas* (orange). Different letters denote signficant differences between groups. Differential abundances were analyzed by ART-ANOVA, FDR*<* 0.05, only non-significant (NS) changes are marked, rest were significant.

PAM1 showed the highest similarity in community composition between samples (Species occurrence/Sorenson’s Index, **Figure 3C**), and had intermediate levels of diversity (OTU diversity, **Figure 3D**) and bacterial load (**Figure 3F, S3C**). Twenty of the 22 most prevalent community members were enriched in incidence and abundance in PAM1 when compared to the other PAMs (ART-ANOVA, FDR*, p*< 0.01, **Figure 3H**, **Table S4**) with five OTUs occurring in >90% of the samples (incidence); *P. melaninogenica* (OTU_3, 97.4%)*, S. mitis* (OTU 11, 99.1%), *V. atypica* (OTU 6, 93.9%)*, V. dispar* (OTU_30, 93%) and *G. adiacens* (OTU_17, 93 %). Contrastingly, two OTUs (*P. aeruginosa*; OTU 1 and *P. fluorescens*; OTU 15) had neither a higher incidence nor a higher abundance in PAM1 (**Figure 3H**, **Table S4**). Thus, PAM1 samples harbor balanced bacterial communities of relatively high similarity composed of the most prevalent bacteria across our dataset. Henceforth, we refer to this PAM as the ‘balanced pneumotype’ (Pneumotype_balanced_).

In contrast to Pneumotype_balanced_, PAM2 and PAM4 harbored lower bacterial diversity (**Figure 3D**) and OTU richness (**Figure 3E**), were dominated by a single community member (**Figure 3G**), and had higher bacterial loads (**Figure 3G**). In these two PAMs, the taxa associated with Pneumotype_balanced_ had a low sample incidence and absolute abundance compared to the other PAMs (**Figure S4A, B**). *S. aureus* (OTU_2), *Corynebacterium* (OTU_24) and *Anaerococcus* (OTU_49) were enriched in PAM2 (ART-ANOVA, FDR*, p*< 0.001, **Figure S4A**), while *Haemophilus* (OTU_78) and *P. aeruginosa* (OTU_1 & OTU_15) dominated PAM4 (ART-ANOVA, FDR*, p*< 0.001, **Figure S4B**, *p*< 0.001). We refer to these as ‘Pneumotype*_Staphylococcus_*’ (PAM2) and ‘Pneumotype*_Pseudomonas_*’ (PAM4), with the major species known to be potential pathogens that proliferate rapidly in lung, under a variety of pathological respiratory conditions (Cohen et al., 2016; Winstanley et al., 2016). Concordantly, the BALF samples assigned to these two pneumotypes were those with the highest bacterial loads.

The fourth cluster identified, PAM3, exhibited the lowest between-sample similarity in species composition (**Figure 3C)**, the highest OTU diversity and richness **(Figure 3D, E)**, and lowest dominance (**Figure 3G**). The samples in this PAM were characterized by considerably low bacterial loads, up to two orders of magnitude below samples in other PAMs (**Figure 3F, Figure S3C**), suggesting a depauperated microbiota that has been associated with dysbiotic physiological states of the gut microbiota (Vandeputte et al., 2017). Consequently, the high OTU richness detected in PAM3 samples is likely due to over-sequencing of rare or transient species, or sequencing artefacts. This is also supported by the fact that the 30 predominant microbiota members were significantly reduced in their incidence and abundance in PAM3 as compared to the other PAMs (ART-ANOVA, FDR*, p*< 0.001**, Figure 3I**). We refer to this PAM as the ‘microbiota-depleted pneumotype’ (Pneumotype_MD_).

Semi-quantitative culture results obtained from matched BALF and bronchial aspirate (BA) on selective media further reinforced the genuine existence of the four pneumotypes (**Figure 3J, S5**, **Methods**). BALF samples from Pneumotype_balanced_ had the highest percentage of matches to the oropharyngeal microbiota, including many of the bacteria that are predominant in this pneumotype (e.g. *Streptococcus* or *Veillonella*). Similarly, culture results of BALF samples with Pneumotype_Staphylococcus_ and Pneumotype_Pseudomonas_ were most frequently positive for *S. aureus/Corynebacterium* spp. and *P. aeruginosa*, respectively, while those obtained for BALF samples with Pneumotype_MD_ were often culture negative (**Figure 3J, S5A**). A similar picture was observed for BA (**Figure 3J, S5B**). Here, however, this sampling site had a higher percentage of positive cultures for oropharyngeal flora compared to BALF, especially for Pneumotype_MD_. This reveals different degrees of segregation between the two sample types, despite the known topographic continuity of microbial communities in the airways (Charlson et al., 2011; Dickson et al., 2015, 2017). Many of the OTUs of Pneumotype_MD_ could not be cultured in our culturomics approach (**Dataset S4, S5**), which together with the low bacterial abundance in corresponding samples, questions their relevance/existence as lung microbiota members. In contrast, most of the major community members characteristic of the other three pneumotypes were represented by isolates in LuMiCol, including the two opportunistic pathogens, *P. aeruginosa* (OTU_1) and *S. aureus* (OTU_2), providing the basis for future experimental work on the predominant strains in the two corresponding pneumotypes. Taken together, we identified four distinct bacterial communities in transplanted lung, which we refer to as pneumotypes, and validated them by semi-quantitative culturing of BALF samples.

### Bacterial pneumotypes are linked to distinct host gene expression patterns

The existence of bacterial pneumotypes with distinctive community composition suggests differences in the microenvironmental conditions of the human lung post-transplant, which could be echoed in other constituents of the lung ecosystem. We compared the median expression levels of 31 host genes belonging to 7 functional categories across the four pneumotypes. These genes are involved in inflammation, immunoregulation, tissue remodeling and detection of bacteria and viruses (**Figure 4A, Methods**). Based on median gene expression, the four pneumotypes showed distinct patterns, with particularly high transcriptional activity in Pneumotype_Staphylococcus_ (**Figure 4A**). To identify the genes with the greatest power to discriminate between the four pneumotypes and to distinguish between samples differing by bacterial counts, we applied a machine learning approach (Random Forest, **Methods**) based on the host gene expression in 234 BALF samples. The Pneumotype_balanced_ was predicted with highest accuracy (92%), followed by Pneumotype_Pseudomonas_ (83.4%) and Pneumotype_MD_ (81.4 %), while no accuracy was achieved for Pneumotype_Staphylococcus_ (**Table S5**). We identified 6 of the 31 genes to have a particularly high predictive power IFNLR1, MRC1, IL10, IL1RN, LY96, IDO (Importance score >10; 99% Confidence Interval, **Figure 4B**). IFNLR1 encodes interferon lambda receptor 1, which is involved in antiviral defence and epithelial barrier integrity (Odendall et al., 2017). This gene was up-regulated in samples with Pneumotype_balanced_ compared to the other three pneumotypes (**Figure 4C**). MRC1 (Mannose Receptor C-Type 1, Geijtenbeek and Gringhuis, 2009) and LY96 (Lymphocyte Antigen 96, Shimazu et al., 1999) encode microbial polysaccharide and lipopolysaccharide recognition proteins, respectively. Compared to samples with Pneumotype_balanced_, these two genes were up-regulated in Pneumotype_Staphylococcus_ and Pneumotype_MD_, and down-regulated in samples with Pneumotype_Pseudomonas_ (**Figure 4D and 4E**). Samples with Pneumotype_balanced_ further differed from those linked to the other three pneumotypes by higher expression of genes involved in immune modulation and peripheral immune tolerance (IL-10/Interleukin 10 and IDO1/Indoleamine 2,3-Dioxygenase 1, **Figure 4F, S6**), and a lower expression of IL1RN (Interleukin 1 Receptor Antagonist, **Figure 4G**), produced as part of the inflammatory response to control the potentially deleterious effects of Interleukin-1 beta (IL-1β) (Arend et al., 1998).

**Figure 4.**
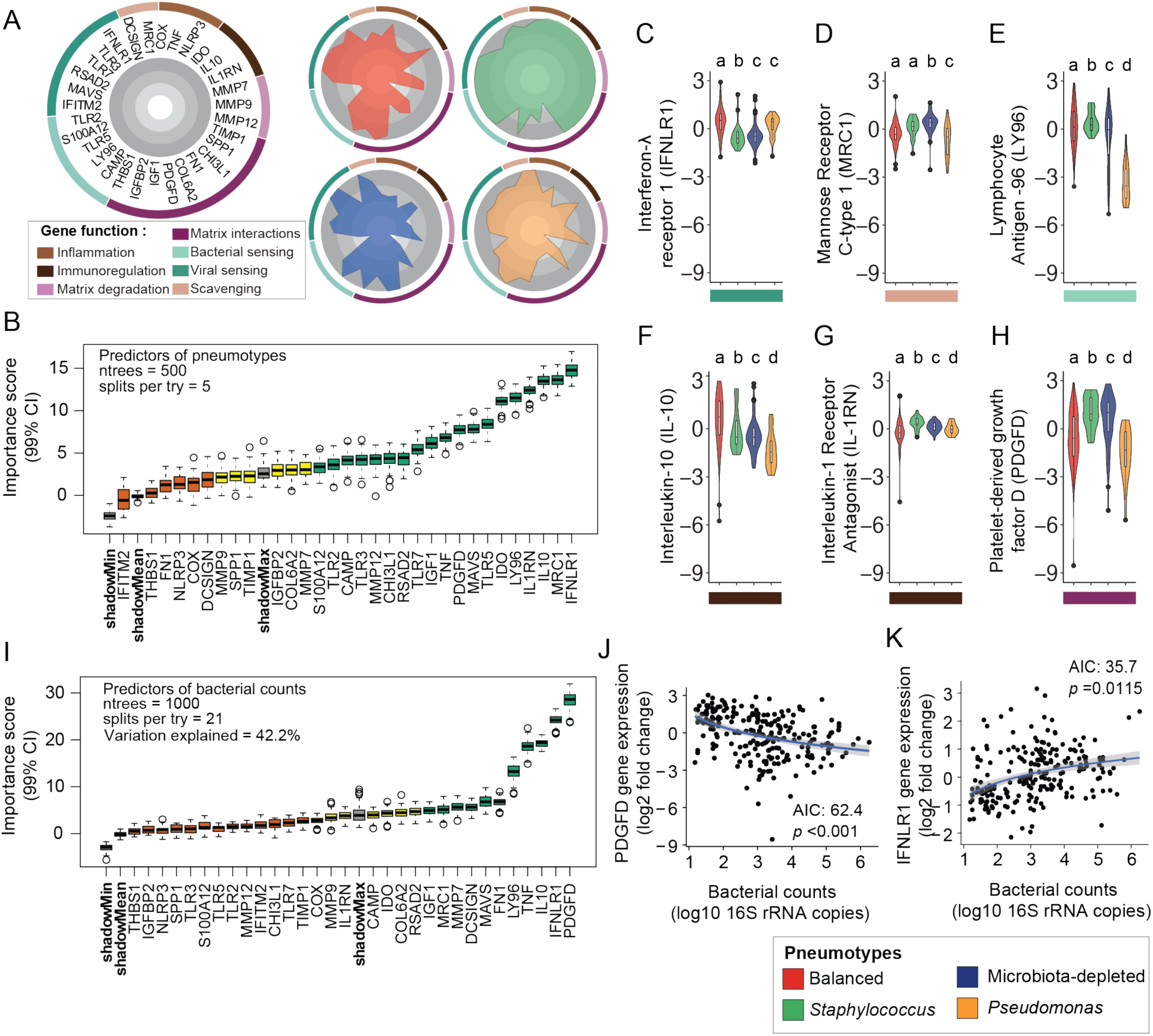
Host gene expression in the lung differs according to pneumotype and bacterial load. (**A**) Radar plots show median-normalized expression of 31 host genes (radial axes) in the cell fraction of all BALF samples (See Methods) split into four pneumotypes. Circular distribution of genes in the plot was colour-coded according to 7 functional categories. Ticks (grey shading) show increase in expression from the inside to outside of circle (**B**) Boxplots show importance (99% Confidence Interval) of host genes as predictors of pneumotypes analysed by Random Forest and Boruta feature selection (See Methods, See also **Table S5**). (**C-H** Violin plots (with inset boxplots) showing log2 expression for 5 of the 6 host genes with Importance scores >10 plus PDGFD across the colour-coded pneumotypes. (**I**) Boxplots show host gene predictors (99% Confidence Interval) of bacterial counts. (**J and K**) Scatter plots show correlation of PDGFD and IFNLR1 expression (log2 fold) with bacterial counts across samples. Boruta provides importance scores to host gene expression predictors and categorizes into Confirmed (Green), Tentative (Yellow) and Rejected (Orange). ntrees = number of decision trees constructed and splits per try = number of random predictors that were sub-sampled at each step and in case of regression provides percent explained variance (See Methods). The most significant model of correlation was selected by stepwise regression (stepAIC; AIC: Akaike Information Criteria) with integrated ANOVA, *p*< 0.05. Differences between pneumotypes were analysed using either one-way ANOVA with Tukey’s post hoc test or Kruskal-Wallis test with Dunn’s post hoc, with different letters denoting significance.

Similarly, we found 5 genes with high discriminating power (**Figure 4I**, importance score > 10) for bacterial counts, of which two were particularly good predictors: PDGFD and IFNLR1. PDGFD encodes the D isoform of platelet-derived growth factor, which promotes the proliferation of cells of mesenchymal origin such as fibroblasts (Simon et al., 2002). Expression of this gene was negatively correlated (**Figure 4J**, AIC 62.4, *p*< 0.001) with bacterial abundance. In contrast, IFNLR1 expression positively correlated with bacterial abundance (**Figure 4K**, AIC 35.7, *p*= 0.0115). Accordingly, PDGFD expression was higher while IFNLR1 expression was lower in Pneumotype_MD_ (**Figure 4C, H**) as compared to the other pneumotypes, suggesting a link between the normal presence of bacteria in the lower respiratory tract and homeostatic levels of tissue remodeling, epithelial barrier integrity and host response to viruses. In summary, these results show that host-specific gene expression markers align with distinct bacterial states, highlighting the existence of complex associations between different lung ecosystem characteristics.

### Anellovirus dynamics is associated with bacterial community and host physiology in lung

The observed links between pneumotypes and antiviral defence prompted us to look into the tripartite interactions between lung bacteria, viruses, and host. To this end, we quantified the load of the three genera of anelloviruses identified in humans (*Alphatorquevirus*, *Betatorquevirus* and *Gammatorquevirus*) across the 234 BALF samples. In accordance with a previous study (Young et al., 2015), we found that the transplanted lung contains high levels of anelloviruses, with *Gammatorquevirus* predominating. Viral loads of the three genera peaked between 1.5 and 6 months after transplantation and decreased at later time points (**Figure 5A**). Anellovirus load varied substantially between pneumotypes. Specifically, the load of *Alphatorquevirus* was highest in samples with Pneumotype_Pseudomonas_ (**Figure 5B**), while that of all three anellovirus genera was lowest in samples with Pneumotype_MD_ (**Figure 5B-D**). This suggested in particular a difference in viral load between Pneumotype_Balanced_ and Pneumotype_MD_, which we confirmed by intra-individual pairwise analysis showing a strong decrease in load when transitioning from Pneumotype_Balanced_ to Pneumotype_MD_ and a corresponding increase for an inverse transition (**Figure 5E, S8**). We further identified 4 human genes: TLR3, IGF1, RSAD2, IFITM2, as important predictors of anellovirus loads in BALF (**Figure 5F, Methods**). Of these, Toll-like Receptor 3 (TLR3) was positively correlated with total viral load (**Figure 5G**, AIC 73.9, *p*<0.001). This is consistent with the low viral load observed with Pneumotype_MD_, where TLR3 was down-regulated (**Figure 5H**). These findings link changes in the lung microbiota composition to changes in viral loads and host gene expression indicating possible implications for allograft outcome.

**Figure 5.**
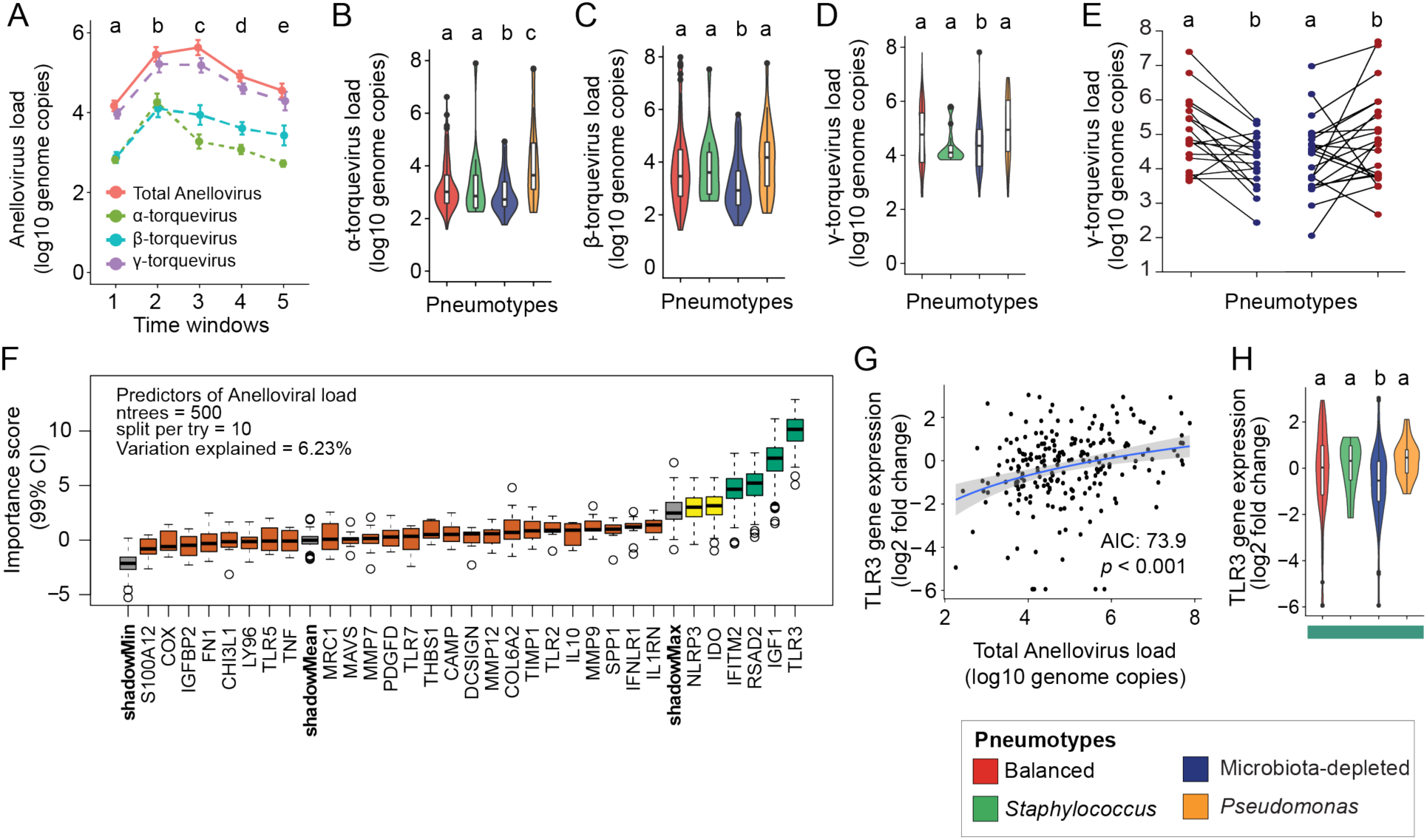
Anellovirus loads differ according to pneumotype and correlate with host physiology in the transplanted lung. (**A**) Line plots show longitudinal progression of Anellovirus load (log 10 pan-Anelloviridae genome copies, salmon pink) and its three major genera: α-torquevirus (Apple green), β-torquevirus (turquoise) and γ-torquevirus (violet) over 5 time windows after transplantation (x-axis). Statistical significance is shown for total viral loads against time windows (one-way ANOVA, *p*< 0.05). (**B-D**) Violin plots (with inset boxplots) show genome copies for individual genera: α-torquevirus, β-torquevirus and γ-torquevirus across pneumotype (plot colors). (**E**) Intra-individual pairwise analysis of γ-torquevirus loads upon transition from Pneumotype_Balanced_ (Red) to Pneumotype_MD_ (Blue) and vice-versa. (**F**) Boxplots show importance of host genes as predictors (99% Confidence Interval) of anellovirus load analysed by Random Forest and Boruta feature selection (For details and gene function categories, See Methods, Figure 4). Scatter **(G)** and Violin **(H)** plots show TLR3 expression correlating with total anellovirus genome copies, and across the four colour-coded pneumotypes, respectively. The most significant model of correlation was selected by stepwise regression (stepAIC; AIC: Akaike Information Criteria) with integrated ANOVA, *p*< 0.05. Differences between pneumotypes were analysed using either one-way ANOVA with Tukey’s post hoc test or Kruskal-Wallis test with Dunn’s post hoc, with different letters denoting significance.

### Pneumotypes are linked to differential risk of post-transplant clinical complication

A large set of clinical data (**Dataset S7**) enabled us to associate differences in bacterial community composition, host gene expression, and anellovirus load to allograft and patient health status. Immunosuppression as well as prophylactic and therapeutic antibiotic usage were anticipated as major confounding factors. However, we found no association between the different pneumotypes and the main immunosuppressive drugs (ANOVA, prednisone; *p*=0.76, tacrolimus; *p*= 0.78) used in our cohort, at the time of BALF sampling (**Figure S7A, B**). In contrast to what has been reported for blood plasma after transplantation (De Vlaminck et al., 2013), we also did not observe a correlation between anellovirus load and immunosuppressive drug levels (Linear regression, *p*=0.91, **Figure S7C, D**, **Methods**). However, we observed a negative relationship between the number of antibiotics administered at the time of BALF sampling and the fraction of samples in Pneumotype_Balanced_ and a positive relationship with the fraction of Pneumotype_MD_ samples (Fisher’s test, *p=* 0.002, **Figure 6A**). These observations thus suggest a link between intensive antibiotic use and a disturbance of the most balanced and compositionally stable lung microbiota profile.

**Figure 6.**
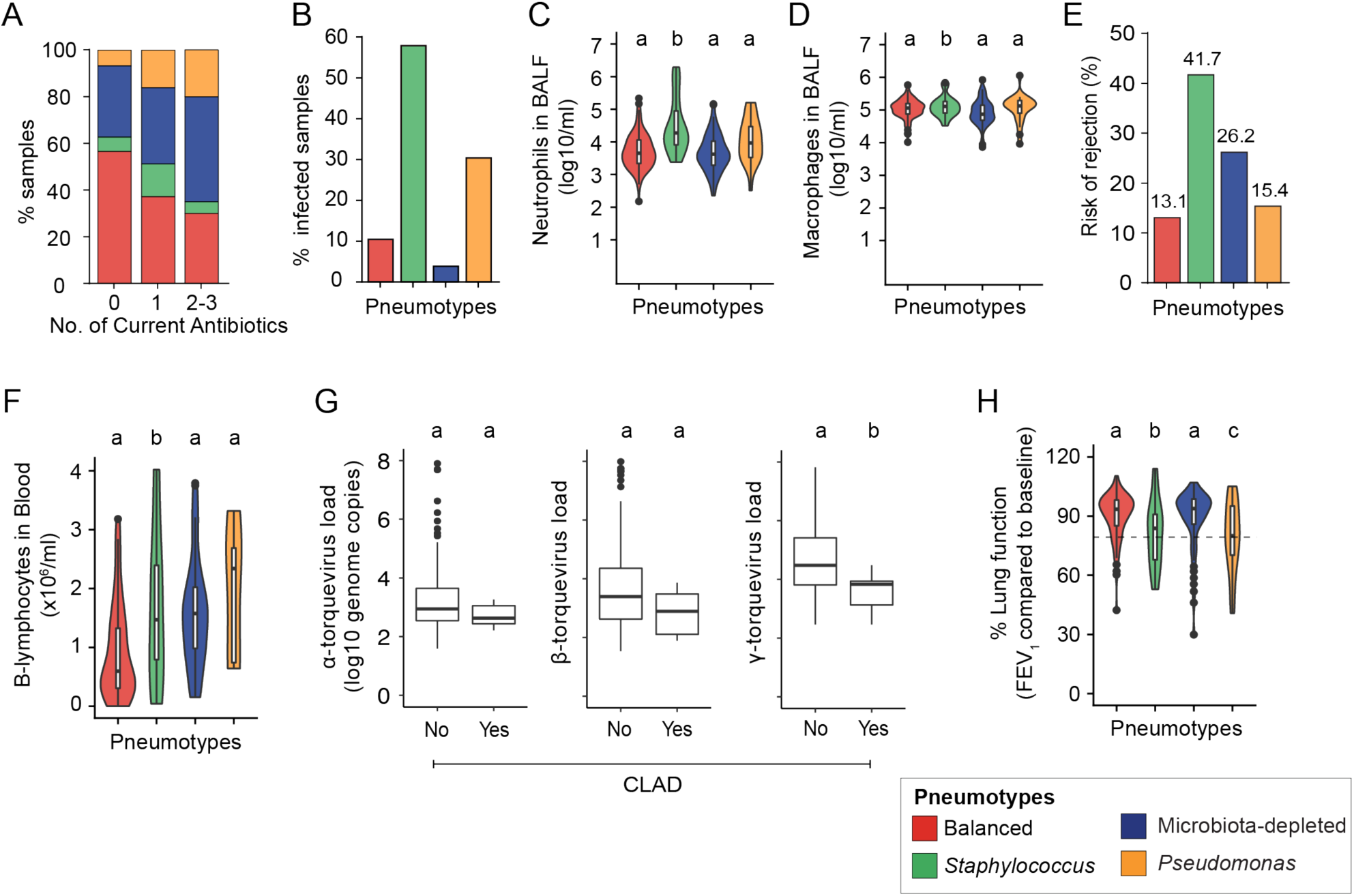
Association of post-transplant pneumotypes with pulmonary environment, local and peripheral host immunity and clinical status. (**A**) Stacked bar plots showing proportion of samples associated with the four pneumotypes relative to the number of antibiotics administered. (**B**) Bar plots show the proportion of infected samples in association with the four pneumotypes. (**C and D**) Violin plots showing numbers of Neutrophils **(C)** and Macrophages **(D)** in lung (log 10 cells per ml BALF) linked to pneumotypes (plot colors). (**E**) Risk of rejection associated with each pneumotype (bar colors) was assessed by the cumulative percentages (%) of samples associated with given conditions (See Methods): Chronic Lung Allograft Dysfunction (CLAD), presence of Donor-specific antibodies (DSA, Mean Fluorescence Intensity > 1000) or Acute cellular rejection (Biopsy score A2) (**F**) Violin plots showing number of B-lymphocytes in the blood associated with the four pneumotypes (plot colors). (**G**) Boxplots show burden of three major anellovirus genera: α-, β- and γ-torquevirus (log10 genome copies) in samples associated with CLAD (Yes or No). **(H)** Violin plot show comparison of lung function (% compared to baseline) measured by Forced Expiratory Volume in 1 second (FEV_1_) across four pneumotypes (plot colors). Differences between pneumotypes were analysed using either one-way ANOVA with Tukey’s post hoc test or Kruskal-Wallis test with Dunn’s post hoc, and difference relative to CLAD was shown by paired Wilcoxon Rank sum test (Mann-Whitney test). Different letters denote significance.

We observed that a clinical diagnosis of infection was rare in the presence of Pneumotype_Balanced_ and Pneumotype_MD_, compared to Pneumotype_Staphylococcus_ and Pneumotype_Pseudomonas_ (Generalized Linear Model, *p*< 0.001 and *p*= 0.016, respectively; **Figure 6B**). This confirms the results of our 16S rRNA gene analysis, which showed that Pneumotype_Staphylococcus_ and Pneumotype_Pseudomonas_ are dominated by the opportunistic pathogens *S. aureus* and *P. aeruginosa,* respectively. It is also consistent with the finding of lower numbers of neutrophils (**Figure 6C**), but not macrophages (**Figure 6D**), in Pneumotype_Balanced_ and Pneumotype_MD_ as compared to Pneumotype_Staphylococcus_ and, to a lesser extent, Pneumotype_Pseudomonas_, emphasizing that pneumotypes are associated with local conditions that differ in terms of recruitment of pro-inflammatory cells.

Lung transplant recipients face risks of allogeneic responses against the graft, notably promoted by clinical infection. Our study did not have the statistical power to dissect the links between pneumotypes and different types of rejection, limited by the number of samples per rejection category in our dataset. Therefore, we grouped 29 samples from 17 patients with either CLAD, acute cellular rejection grade ≥2, or the presence of donor-specific antibodies (mean fluorescence intensity >1000), as these all indicate a suboptimal control of host immune competence and thus an increased probability of allograft injury (**Figure 6E, See Methods for clinical definitions**). The majority of these samples were associated with Pneumotype_Staphylococcus_ (41.7%) and Pneumotype_MD_ (26.2%), followed by Pneumotype_Pseudomonas_ (15.4%) and Pneumotype_Balanced_ (13.1%), suggesting that this latter microbiota profile is associated with a lower risk of clinical complications. This was further corroborated by the count of circulating B lymphocytes in peripheral blood, suggesting more active humoral immunity in the presence of Pneumotype_MD_, and to a lesser extent for Pneumotype_Staphylococcus_ and Pneumotype_Pseudomonas_, compared to Pneumotype_Balanced_ (Tukeys test, *p*= 0.027, *p*= 0.26 and *p*= 0.12 respectively; **Figure 6F**). In addition to bacterial composition, anelloviruses were also linked to CLAD through a lower load of *Gammatorquevirus* (Wilcox test, *p*= 0.007, **Figure 6G**), while no significant association was observed with *Alphatorquevirus* or *Betatorquevirus* (Wilcox test, *p*= 0.15 and *p*= 0.09 respectively, **Figure 6G**).

Finally, we used the measurement of ‘Forced Expiratory Volume in one second’ (FEV1) to search associations between lung ecology and pulmonary function testing. This assessment was made irrespective of the diagnosis of CLAD, which requires an irreversible drop in FEV1 below 80% of the baseline value, with prior exclusion of alternative confounding diagnosis **(See Methods).** Pneumotype_Staphylococcus_ and Pneumotype_Pseudomonas_ were associated with lower FEV1 values overall, with a frequent substantial decline below 80% predicted (Dunn’s test, *p=* 0.03, **Figure 6H**), while Pneumotype_Balanced_, along with Pneumotype_MD_, was linked to preserved lung function.

### Pneumotype_Balanced_ shows the highest temporal stability and resilience in the transplanted lung

Taking advantage of the longitudinal sampling, we explored the dynamics of pneumotypes after transplantation. We analyzed transitions between pneumotypes in up to eight BALF samples per transplant, collected within five consecutive time windows (**Figure 7A**). There was no significant difference in the distribution of pneumotypes across the different time windows (Chi-square test, *p*= 0.60). Although most BALF samples were associated with Pneumotype_Balanced_, transitions between two different microbiota profiles occurred for about half of all consecutive sample pairs (**Figure S9A**). The transition dynamics were explained by Markov chain properties, i.e. the pneumotype of a given sample only depends on the state of the previous sample in the chain (Chi-square test, *p*= 0.33, **Figure 7B**). The transitions were irreducible, aperiodic and recurrent, and none of the pneumotypes behaved as an absorbing state **(See Methods)**. Pneumotype_Balanced_ exhibited the greatest stability, with the highest probability of recurrence (63%), fitting the Markov probabilities, followed by Pneumotype_MD_ (42%; Figure 7B). In addition, the large fraction of transitions towards Pneumotype_Balanced_ between the first four time windows indicated a substantial resilience capacity for this profile. Accordingly, the transitions between Pneumotype*_Staphylococcus_*, Pneumotype*_Pseudomonas_* and Pneumotype_Balanced_ occurred mainly in the direction of this latter profile (**Figure 7B**), while in contrast to model prediction, Pneumotype*_Staphylococcus_* and Pneumotype*_Pseudomonas_* appeared to be virtually disconnected.

**Figure 7.**
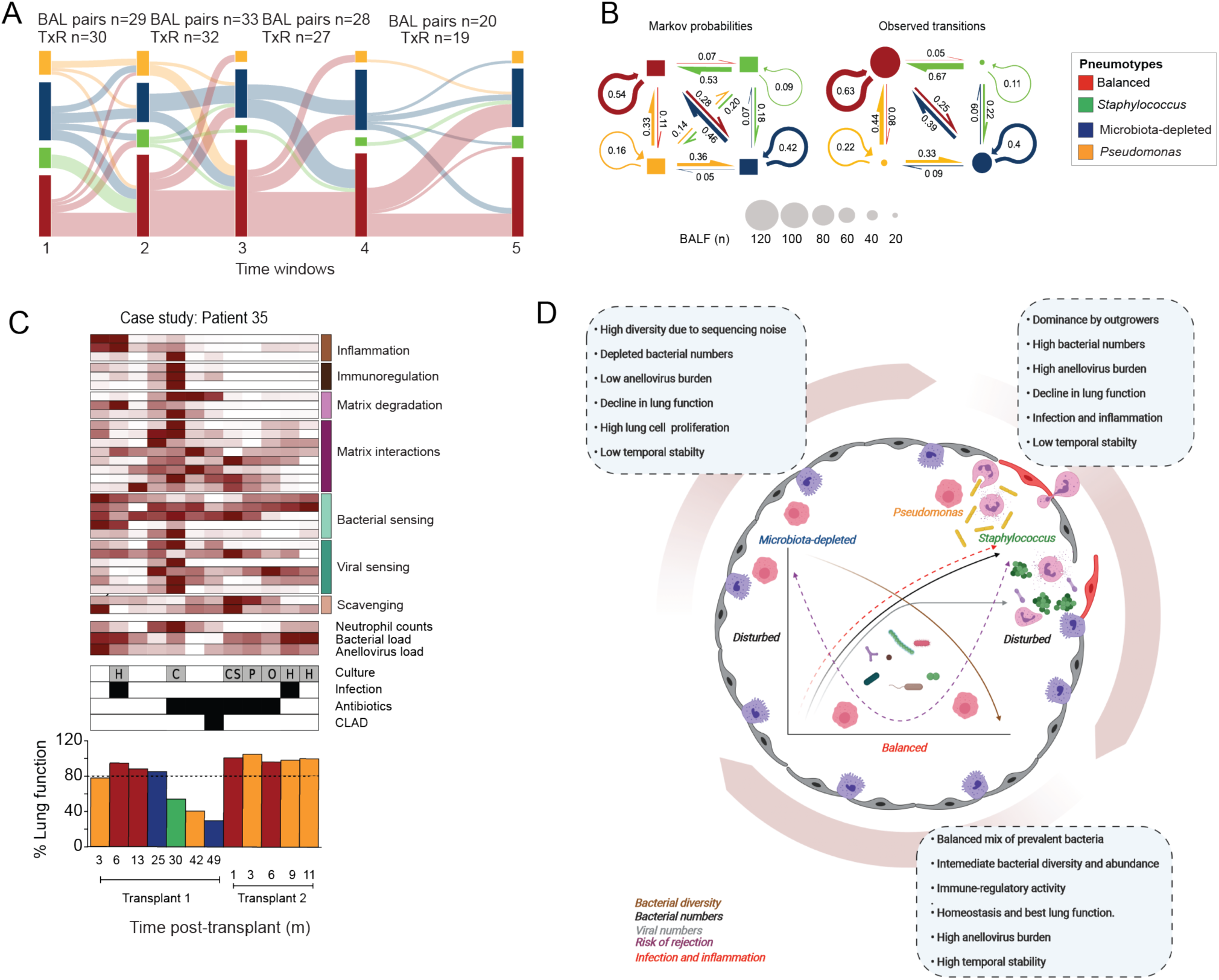
Longitudinal analysis of lung microbiota post-transplant and dynamics of pneumotype transitions. **(A)** Sankey diagram showing transition of paired samples between pneumotypes (colors) across 5 Time Windows. **(B)** Markov chain model (See Methods) fitted to the observed pneumotype transitions (n = size of circle). Model was initiated with equal probabilities for each pneumotype (0.25, 100 bootstraps, left panel) and given transition matrix. Pneumotypes are represented by colored arrows/boxes and direction of a transition is indicated by a colored arrow of a thickness denoting the probability. **(C)** A patient case study showing transition of pneumotypes with clinical characteristics across two transplantation events. Heatmap shows host gene expression with functional categories (See also Figure 4A, right vertical colored bars), neutrophil counts, bacterial and anellovirus loads in BALF across time and pneumotypes. Taxa obtained in routine clinical culture were abbreviated with letters. Samples positive for infection, ongoing antibiotic treatment or CLAD (black boxes) are presented above bar plots showing % lung function (See also Figure 6G), across transplantation events and time post-transplantation (months) and pneumotypes (bar colour). **(D)** Scheme of bimodal disruption in lung ecosystem (colored arrows in a x-y plot) leading either to (i) a microbiota-depleted pneumotype with ambigous bacterial diversity (brown), low counts of bacteria (black) and viruses (grey), high lung cellular proliferation and chronic decline in lung function leading to rejection (purple), or *ii*) pneumotypes dominated by opportunists (*Staphylococcus* and *Pseudomonas*) with loss in bacterial diversity, high infection rate and inflammation (red), acute decline in lung function and rejection. Best-case scenario is defined by a middle ground with a balanced pneumotype consisting of the most prevalent bacteria in a homogenous composition with intermediate bacterial diversity, bacterial and viral abundance, high immune-modulatory activity and best preserved lung function.

Finally, we illustrate the relationship between the temporal dynamics of pneumotypes with clinical outcomes, using a case study (**Figure 7C**). Patient 35 diagnosed with pulmonary fibrosis received two transplants, providing 12 serial samples and presenting each of the four pneumotypes (**Figure 7C - bottom histogram**). Disruption of the Pneumotype_Balanced_ occurred from month 25, followed by transition to Pneumotype_Staphylococcus_ at month 30. This was accompanied by a positive culture for *Corynebacterium* spp. in line with our enrichment analysis and clinical culture tests (**Figure S4, Figure 3J**), increased BAL neutrophilia, and a concurrent increase in host immune gene expression (**Figure 7C - heatmap**). Thereafter, the patient was repeatedly exposed to antibiotics, and respiratory function started declining irreversibly leading to the diagnosis of CLAD at month 49 with Pneumotype_MD_. Overall decrease in lung bacterial and anellovirus loads in the lung, suggested an increasing selection pressure on microbes most likely due to a combination of antibiotic treatment and poorly controlled host immune competence. The second transplant at month 50 was linked to a re-establishment of Pneumotype_Balanced_, which aligned with preserved lung function, intermediate loads of lung bacteria and anelloviruses, decrease in neutrophil counts and change in host immune gene expression. However, a transition to Pneumotype_Pseudomonas_ was observed later until the end of sampling, with increased bacterial counts but no decrease in lung function (**Figure 7C - barplot**). Taken together, these observations highlight the potential of integrating pneumotype with clinical and molecular data, with the primary goal of tracking disruption of Pneumotype_Balanced_ beneficial to clinical stability.

## Discussion

In the current study, we capitalized on the availability of 234 longitudinal BALF samples from 64 lung transplant patients. We combined culture-dependent and -independent approaches to characterize the composition of the human lung microbiota, to obtain representative cultured isolates and test their growth requirements, to assess the temporal dynamics of these communities in the lung, and ultimately to establish links with the host health status. In summary, our results show that the lung microbiota post-transplant is highly dynamic with a few predominant community members, many of which can be cultured, under different physiological conditions. We find that the lung microbiota post-transplant can be categorized into four compositional states, ‘pneumotypes’, based on distinctive bacterial community features. These pneumotypes have different temporal dynamics and bridge the gap between lung bacteria, anellovirus loads, host gene expression, and the physiological and immunological state of transplant recipients. Altogether, these results provide important advances in our understanding of lung bacterial communities, their clinical significance, and the experimental tractability of major lung bacteria.

Our analyses show that the human lung microbiota post-transplant predominantly consists of oropharyngeal taxa similar to the microbiota of healthy lungs (Erb-Downward et al., 2011; Segal et al., 2013; Venkataraman et al., 2015). Hence, the presented results are not only relevant in the context of lung transplantation, but also provide general insights into the microbial ecology of the lower respiratory tract. Besides the high variability in taxonomic composition, we find that the total bacterial biomass in the lung can considerably vary between samples. Such quantitative differences in lung microbiota composition have also been found in previous studies in the lung. We find that a relatively small number of OTUs accounted for a large part of the total bacterial biomass detected across all samples (19 OTUs contributing >75% of the biomass), despite the detection of more than 7000 OTUs. These included not only prevalent oropharyngeal taxa but also potential pathogens that outgrew in a few samples. These findings, and the fact that a key characteristic of Pneumotype_MD_ is its association with low bacterial biomass, highlight the importance of considering absolute bacterial counts instead of relying only on proportional data in microbiome studies (Kešnerová et al., 2020; Vandeputte et al., 2017). This is further evidenced by the fact that bacterial biomass can be predicted by host gene expression.

In addition to considering total bacterial biomass, demonstrating the viability of bacteria taxa detected by sequencing can provide further insights about their biological relevance of specific community members. Previous studies have shown that bacteria from the human lung can be cultured with the majority growing under oxic conditions (Cummings et al., 2020; Venkataraman et al., 2015; Whelan et al., 2020). Our large-scale culturomics approach, which included a wide array of culturing conditions, substantially expands the availability of bacterial isolates from the human lung and provides important new insights about their phylogenetic diversity, physiological preferences, and metabolic potential. For instance, we show that many isolates, including prevalent community members, preferred to grow under anaerobic or microaerophilic conditions, suggesting the presence of regions with low oxygen concentration in the deep lung. Also, the culturing allowed us to phenotypically describe specific isolates in more detail and identify closely related pathogenic and non-pathogenic strains of *Streptococcus*, which otherwise could not have been discriminated based on amplicon sequencing alone. Notably, the genus *Streptococcus* had the highest genetic, metabolic, and phenotypic diversity among all isolates, which may explain its presence throughout the human respiratory tract including sites with very different physicochemical properties (Dickson et al., 2017). We acknowledge that the presented culture collection is not exhaustive and several major community members have not yet been isolated. We believe that this is most likely due to the high variability of the lung microbiota and the fact that we have cultured a relatively small number of BALF samples, rather than the inability of some community members to grow *in vitro* or non-viability in the lungs.

The high variability in community composition and bacterial load between BALF samples may suggest that the human lung microbiota is highly erratic. However, our unsupervised machine learning approach identified four compositional states, Pneumotype_Balanced_, Pneumotype_MD_, Pneumotype*_Staphylococcus_* and Pneumotype*_Pseudomonas_*, with distinct community properties. In a previous study on the lung microbiota of healthy individuals, a similar approach was used which resulted in the identification of two pneumotypes (Segal et al., 2013). Strikingly, one of these previously identified pneumotypes was enriched in supraglottic taxa, i.e. mainly *Prevotella, Streptococcus*, and *Veillonella* resembling the Pneumotype_Balanced_ from our study. The other pneumotype described by Segal et al. had similar characteristics as Pneumotype_MD_ i.e. very low bacterial counts and a highly variable taxonomic composition. As with Pneumotype_MD_ in our study, many of the taxa in this other pneumotype were considered to represent contaminations (or so-called background taxa). In contrast, Pneumotype*_Staphylococcus_* and Pneumotype*_Pseudomonas_* were not detected in this previous study probably because it was based on smaller cohort size and exclusively included samples from healthy individuals. Interestingly, *Staphylococcus*, the major community member of Pneumotype_Staphylococcus_, has been shown to dominate in neonatal lower airways, indicating potential early adaptation to human lung (Pattaroni et al., 2018). Together, these studies provide independent evidence for the existence of distinct compositional states of the human lung microbiota in different contexts. Moreover, the fact that the four pneumotypes are linked to differences in host gene expression, bacterial and anellovirus loads, and allograft function and health state highlights their relevance and suggests the existence of distinct ecological conditions in the lower respiratory tract, which are further discussed in the following sections.

We propose that Pneumotype_Balanced_ is primarily associated with lung homeostasis, because it is characterized by a diverse bacterial community, with a moderate bacterial and viral load, and shows a human gene expression profile leaning towards immune modulation and peripheral immune tolerance. A striking characteristic of Pneumotype_Balanced_ was also the clear association with high expression of Interferon-λ receptor 1 (IFNLR1), which suggests a link between the bacterial community and the maintenance of the epithelial barrier integrity (Odendall et al., 2017) and antiviral defense (Broggi et al., 2020). Moreover, Pneumotype_Balanced_ showed down-regulation of Interleukin-1 receptor antagonist (IL1RN), produced in response to pro-inflammatory cytokines (Arend et al., 1998) and up-regulation of Interleukin-10 (IL-10), a tolerogenic cytokine (Ng et al., 2013). This along with the previously reported association with Th17 immune response (Segal et al., 2016), indicates a possible role of Pneumotype_Balanced_ in development of regulatory T cell and the maintenance of immune surveillance, as seen in case of gut bacteria (Atarashi et al., 2013; Ivanov et al., 2008). In line with this, individuals with Pneumotype_Balanced_ had the lowest risk of clinical complications at the time of sampling. Moreover, transitions from Pneumotype_Balanced_ to other pneumotypes were the least frequently observed. Overall, these observations corroborate the steady state associated with this pneumotype, suggesting that it is indicative of lung health and clinical stability after transplantation.

We show that the gene expression of a hallmark of M2-like macrophages, Mannose receptor C-type 1 (MRC1) (Geijtenbeek and Gringhuis, 2009) and an important contributor to airway remodeling, Platelet-derived growth factor-D (Simon et al., 2002), were increased in Pneumotype_MD_. The low microbial load and the associated loss of a steady-state inflammatory level could be the underlying cause for the increased expression of these genes resulting in unrestrained host cell proliferation and increased deposition of extracellular matrix, as observed in CLAD (Verleden et al., 2020). Another striking feature of this pneumotype was the low expression of TLR3, a host gene involved in virus detection, which was consistent with the low loads of anelloviruses observed in the lung in the presence of this pneumotype. Virtually all lung transplant recipients carry anelloviruses, mainly in the plasma but also in lung, with viral load fluctuating over time (De Vlaminck et al., 2013; Young et al., 2015). Previous reports have shown that anellovirus counts in plasma are associated with host immunecompetence, infection and alloimmune rejection (Abbas et al., 2017; Blatter et al., 2018; Görzer et al., 2017; Jaksch et al., 2018; Segura-Wang et al., 2018; De Vlaminck et al., 2013), indicating a stronger selective pressure imposed by the host immune system in Pneumotype_MD_ on viruses and bacteria in the lung. This was confirmed by low risk of infections and substantial risk of poorly controlled immune activity, which was evident from the high number of circulating B lymphocytes and either donor-specific antibodies, acute cellular rejection or CLAD.

A strongly contrasting pattern was observed for samples with either Pneumotype_Staphylococcus_ or Pneumotype_Pseudomonas_, tightly bound to an inflammatory background. Here, viral and bacterial loads were increased relative to samples with Pneumotype_Balanced_ and Pneumotype_MD_. This was accompanied by a higher risk of infection and a consistent recruitment of neutrophils into the lung, ultimately leading to impaired pulmonary function. Notably, Pneumotype*_Pseudomonas_* was associated with low expression of Lymphocyte antigen (LY) 96 / Myeloid Differentiation protein (MD2), an essential component of the human TLR4 complex (Shimazu 1999). Although it is tempting to associate the importance of *P. aeruginosa* in this pneumotype with a lack of engagement of the TLR4 pathway in the host (Awasthi et al., 2019; Faure et al., 2004), we cannot conclude about a causal link. In line with evidence that infection activates alloimmune responses (Chong and Alegre, 2014), samples with either Pneumotype_Staphylococcus_ or Pneumotype_Pseudomonas_ were also associated with a significant risk of poorly controlled immune activity and rejection.

Our study lacked sufficient statistical power required to explore the links between pneumotypes and different types of rejection (acute cellular rejection, antibody-mediated rejection, CLAD). However, grouping these samples allowed us to associate the Pneumotype_Balanced_ with the lowest risk of poorly controlled immune activity. Furthermore, we could not assign causality to the observed links between the different constituents of the lung ecosystem. This was due to both the non-interventional nature of our approach and the multiplicity of confounding factors and their variability across the cohort. In particular, the underlying therapeutic treatments were expected to significantly modulate lung ecology, in addition to the effects due to infection and alloimmune response. This was illustrated by the observed link between Pneumotype_Balanced_ and samples collected in the absence of ongoing antibiotic treatment, as opposed to the association between pneumotypes with disrupted bacterial communities and ongoing antibiotic therapy. Finally, follow-up studies are required to extend the knowledge gained by our single-site BAL sampling, which would not capture potential variability in ecological conditions across different regions of the lung (Jorth et al., 2015) or between the lung and the upper respiratory tract (Dickson et al., 2015; Simon-Soro et al., 2019).

In conclusion, our work provides a foundation for understanding the need for a balanced lung ecosystem along the bacterial community-viruses-host physiology axis, to maintain respiratory function and health. Overall, we propose that the four pneumotypes seem to follow the “Anna Karenina principle”, where healthy communities vary little around a stable state, while perturbed communities in dysbiotic individuals are much more variable with unstable states (Zaneveld et al., 2017). We propose that the integration of multi-omics data analyzed using ecological principles will assist in the management and follow-up of lung transplant recipients, particularly with respect to CLAD prediction and supportive interventions. An important next step will be to establish causal links between lung ecology and allograft health by identifying the microbiota and host-related factors underlying pneumotype transitions. To this end, our strain collection LuMiCol provides a highly valuable resource that will serve as a foundation for future experimental studies using animal or cell culture models.

## Methods

### Statistical analysis and Software used

Various statistical approaches and tests were used depending on methods, as detailed in the appropriate sections. Significant differences between groups are denoted by letters. All analyses were performed on R version 3.5.2, python v 2.6 and bash on macOS Mojave v 10.14.6.

Citations were included for all softwares used except for packages available via CRAN Repository and tools that are available via downloads from public database. For sequencing quality control and curation FastQC; https://www.bioinformatics.babraham.ac.uk/projects/fastqc/ and FASTX-Toolkit; http://hannonlab.cshl.edu/fastx_toolkit/index.html were used. A custom pipeline for sequencing analysis was build using QIIME v1.9 (Caporaso et al., 2010), vsearch v 2.3.4 (Rognes et al., 2016), ampvis2 v 2.3.2 (Andersen et al., 2018), phyloseq 1.26.1 (McMurdie and Holmes, 2013) and vegan package version 2.5-6. Alignment and taxonomic classification were obtained using SINA aligner; https://www.arb-silva.de/aligner/ was used on the local computer. Phylogeny was performed by FastTree v 2.1.10 and visualized using iTOL; https://itol.embl.de. K-medoid-based unsupervised machine learning was performed on Genocrunch; www.genocrunch.epfl.ch. Differential abundances was tested using ART-ANOVA from ARTool package version 0.10.7. Machine learning classification and regressions were performed *randomForest* package and its wrapper algorithm *Boruta* for feature selection.

LuMiCol isolate sequences were curated using Geneious Software v 10.2.6., New Zealand. All graphical illustrations were made using BioRender Web Application, Canada.

### Data and code availability

We have deposited the raw data from all samples used in the study to Short Read Archive, NCBI under the BioProject PRJNA632552 and BioSample accession SAMN14911405. All Datasets and codes are available for access on the cloud drive below. Details of Datasets and Tables are mentioned in the Supplementary Data summary.

https://drive.switch.ch/index.php/s/hch0EoA5QyjBPR8

**Das_et_al_2020_analysis_pipeline_1:** Sequencing analysis pipeline using python (QIIME) and bash (vsearch, FastXToolkit, SINA, FastTree). Code from raw data processing, merging cultured sequences (LuMiCol), OTU picking and phylogeny.

**Das_et_al_2020_analysis_pipeline_2**: R markdown with BALF community analysis with starting OTUs from pipeline 1 with phyloseq, ampvis2 and vegan. Random Forest algorithms, Markov chain analysis and all statistical analysis and visualization plots.

**All_BAL_samples_raw_fastqc:** FastQC reports for raw sequencing data after merging all samples.

**All_BAL_samples_processed_fastqc:** FastQC reports for trimmed and curated merged data.

### Study Population, Sampling and Ethics Statement

#### Study design

In this prospective longitudinal study, we used a cohort of 64 consecutive lung transplant recipients from our center. We collected 234 BALF samples (n=1-12 per recipient, mean 3.7) between 2 weeks and 49 months post-transplantation, during routine surveillance or clinically indicated bronchoscopies, from October 2012 to May 2018. Details on BALF collection and processing are provided below.

#### Ethics statement

The study was approved by the local ethics committee (“Commission cantonale (VD) d’éthique de la recherche sur l’être humain – CER-VD”, protocol number 2018-01818), and all subjects gave written informed consent. Samples were anonymized according to local ethics committee requirements.

#### Patient sample collection

Patients underwent transoral bronchoscopy. For BALF collection, the bronchoscope was wedged either in the middle lobe or lingula of the allograft and 100-150 ml of normal saline were instilled in 50 mL aliquots that were pooled. BALF recovery was measured and the sample was submitted to cell differential determination according to routine clinical procedures. Two fractions of 3 ml were stored at 4°C and centrifuged within 3 h at either 2,000 or 14,000 x g for 10 min, for future isolation of BALF cellular RNA and total DNA, respectively. Pellets were snap frozen, either after cell lysis in RLT buffer (Qiagen, Hilden, Germany) to preserve RNA integrity, or directly, and were stored at minus 80°C until further processing. A negative control obtained upon washing a ready-to-use endoscope with sterile saline was prepared following the same procedure.

### Bacterial culturomics and establishment of Lung Microbita culture Collection (LuMiCol)

#### BALF cultivation and archiving

A volume of 100 microliters of BALF was spread per plate of 15 different media (Table 5) within 2 to 3 hours following endoscopy. The plates were then immediately incubated at the desired combination of oxygen and temperature conditions; aerobic (AE), microaerobic (MI; O_2_: 17%, CO_2_: 5%, Relative Humidity: 85%) and anaerobic (AN; H_2_: 8%, N_2_: 72%, O2: 40 ppm, CO_2_: 20%) at a temperature range between 35-37°C (**Dataset S4**). Plates were incubated between 1-5 days. Bacteria were collected from plates by adding RPMI 1640 liquid medium supplemented with 15% Glycerol and scraping using a Drigalski spatula and finally transferred into 96-well plates. Plates were made in triplicates for back up stocks. Each isolate was given a plate identifier (plate number - Px and well number - A1-H12) and a unique isolate code made with a combination of sample number, oxygen condition (AN/MI/AE), Media used and isolate number (**Dataset S4** and **S5**).

#### Genotyping of bacteria

Genotyping and species determination were based on PCR amplification of either Universal 16S rRNA gene (V1-V5 region) or specific marker genes, respectively. *Staphylococcus aureus* was identified by the presence of *nuc* gene encoding staphylococcal thermonuclease. (**Dataset S7**. The sequences were aligned using two well curated databases containing high quality 16S rRNA sequences to resolve species: SILVA SSU rRNA database and wherever SILVA failed to provide species identification we used the extended Human Oral Microbiome Database; eHOMD, http://www.homd.org. Phylogeny was performed by FastTree v 2.1.10 and visualized using iTOL.

#### Bacterial growth determination by optical density

Undefined rich media was represented by Todd-Hewitt (CM0189, Oxoid, UK) supplemented with yeast extract (0.5g/L, LP0021, Oxoid, UK). RPMI 1640 medium with (11875085, ThermoScientific, USA) and without Glucose (11879020, ThermoScientific, USA) represented low complexity defined media. RPMI1640 without Glucose was chosen as a proxy for lung deep lung fluids since it contains free amino acids, physiological salts, Glutathione and no Glucose, which are properties similar to lung epithelial lining(Evans et al., 2014).

One representative of each 47 phylotypes was revived on its individual isolation media (Table 5), and bacterial biomass was scraped off the plates using 1X PBS. Bacterial suspension was diluted into 200 μL medium of appropriate media in 96-well non-tissue culture treated transparent flat-bottom plates (CytoOne^®^, CC7672-7596, Starlab, Germany). The plates were then immediately incubated at the desired combination of oxygen and temperature conditions; aerobic (AE, 37°C), microaerobic (MI; O_2_: 17%, CO_2_: 5%, Relative Humidity: 85%, 37°C) and anaerobic (AN; H_2_: 8%, N_2_: 72%, O2: 40 ppm, CO_2_: 20%, 34°C) (**Dataset S4**). Optical density was measured at 600 nm using a BioTeK Synergy H1 Hybrid Multi-Mode Reader starting from time Day 0 (0 minutes) and everyday (24 hours) up to Day 3 (72 hours). Growth at each time point was calculated by the change in optical density from Day 0 (ΔOD). The experiment was repeated three times and the median ΔOD for each day was used to create a heatmap.

### Species identification by phenotypic assays

To differentiate staphylococcal and streptococcal species, primarily *S. aureus* from other *Staphylococci, and* Viridans *Streptococcus from Pneumococcus,* bacteria were screened for multiple phenotypes. As controls, *Staphylococcous aureus* ATCC 25904, *Streptococcus pneumoniae* strain D39; NCTC 7466 (pneumococcus control) and *Streptococcus mitis* NTCC10712 (viridans *Streptococcus* control, provided kindly by the group of Dr. Jan-Willem Veening, Lausanne, Switzerland) were used. For general overnight culture, *Streptococci* were grown in Todd-Hewitt (CM0189, Oxoid, UK) supplemented with yeast extract (0.5g/L, LP0021, Oxoid, UK) at 37°C with 5% CO_2_, 85% Relative Humidity and *Staphylococci* were grown in Tryptic Soy Agar (CM0131B, Oxoid, UK) at 37°C.

#### Hemolysis detection on semi-solid agar

For detection of hemolysis, bacteria were grown on Columbia agar (CM0331B, Oxoid, UK) supplemented with 5% Defibrinated Sheep Blood (SR0051E, Oxoid, UK) and incubated at 37°C under aerobic conditions or with 5% CO_2_, 85% Relative Humidity and lysis of blood was observed after 24 hours. After which complete hemolysis (beta-hemolysis) can be observed if any and then the plates were transferred to 4°C for partial hemolysis alpha-hemolysis) to be more prominent.

#### High salt growth and Mannitol fermentation test for Staphylococci

The ability of *Staphylococci* to grow on high salt and ferment Mannitol was tested by cultivation on Mannitol Salt Agar (MSA, 7.5% Sodium Chloride and D-Mannitol, CM0085B, Oxoid, UK) and incubation at 37°C under aerobic conditions. This resulted in few combinations: Growth or no growth in MSA, growth in MSA but no fermentation of Mannitol, growth in MSA and also fermentation of Mannitol (designated by the conversion pink Phenol red to yellow color).

#### DNase activity assay

Staphylococcal Thermonuclease activity was tested by growing *Staphylococci* on DNase agar (CM032, Oxoid, UK), as previously described (Fusillo and Weiss, 1959). Briefly, bacteria grown overnight on and a single colony was streaked using a disposable plastic inoculation loop across in a straight line at the center of the agar plate. Plates were incubated at 37°C under aerobic conditions for 24 hours, before flooding with 1N HCl. After a dwell time of 30 seconds, acid was drained out and a halo around the bacterial biomass indicated a positive result for DNase activity.

#### Optochin resistance test

For differentiating between viridans Streptococci from *Streptococcus pneumoniae,* an Optochin resistance test was performed (Bowers and Jeffries, 1955). Briefly, *Streptococci* were spreaded throroughly on Columbia agar (CM0331B, Oxoid, UK) supplemented with 5% Defibrinated Sheep Blood (SR0051E, Oxoid, UK) with a cotton swab before placing Optochin disks (74042, Sigma-Aldrich, Germany) on the center of the plates and incubated at 37°C under microaerophillic conditions i.e. 5% CO_2_ for 24 hours. Inhibition zones were observed the next day for *S. pneumoniae* but not in case of *S. mitis*.

### BALF microbiota community analysis

#### Bacterial 16S rRNA Amplicon sequencing

The 16S content of BALF DNA was characterized either by quantitative PCR using previously reported primers specific to pan bacteria (**See Dataset S7**, which includes references), and Illumina MiSeq sequencing using primers targeting the V1-V2 region was performed as previously described (**See Dataset S7**). Briefly, the V1-V2 region was amplified with barcoded primers and then sequenced on the Illumina MiSeq platform using paired-end chemistry, generating 250 x 2 read lengths.

#### Data processing and OTU picking

Data curation and analysis was performed using a custom pipeline (See Code availability). The major packages used are described in the Statistical analysis and Software section. Primers were removed and reads were joined fastq-join with a minimum overlap of 10 base pairs, demultiplexed and quality filtered (PHRED score Q> 28 in 75% of read length). Sequence quality was assessed using FastQC and first 75 bases were trimmed using fastx_trimmer. Both raw (All_BAL_samples_raw_fastqc) and processed (All_BAL_samples_processed_fastqc) sequence quality analysis are available for open access online.

Singletons were removed using vsearch. Prior to OTU picking, taxa were clustered into centroids with >98% identity, chimeras removed and the data was mapped to the centroids with >97% coerced into a single OTU. The sequences obtained after OTU picking were further used for alignment and taxonomy using SINA aligner using SILVA SSURef_NR99_132_SILVA_13_12_17 release as reference database. Phylogeny was performed by FastTree v 2.1.10 and visualized using iTOL.

In conventional practice, low abundance samples are excluded but we elected to retain them if sequencing was successful (≥10^4^ reads). However, as this increases the risk of obtaining spurious taxa, we removed OTUs with ambiguous taxa i.e. NA and any samples that had less than 10^4^ reads and no information on bacterial abundance. Due to low biomass, it was important that we analyzed the negative controls, which included Bronchoscope pre-wash, DNA extraction reagents and no-template PCR reaction. We found that negative control samples contained 1015 OTUs, including those from family *Enterobacteriaceae* and genera *Limnohabitans, Fodinicola, Staphylococcus, Flavobacterium, Cutibacterium, Acidovorax, Tepidimonas* and *Variovorax.* After quality filtering and normalization, we ended up with 7164 OTUs at 97% identity in 16S rRNA gene. These OTUs belong to 37 phyla with the most abundant phyla being *Bacteroidetes, Firmicutes, Proteobacteria* and *Actinobacteria*.

#### Extraction of OTU-isolate match pairs

In order to identify, exact OTU-isolate match pairs a combinatorial hybrid sequence pipeline to merge 215 high quality 16S rRNA sequences obtained by Sanger sequencing for LuMiCol isolates (phred score: Q30 > 90%) with the 16S amplicon sequencing dataset and performed OTU picking and taxonomic identification for the retrieval of OTU-isolate matching pairs. This was possible due the common forward primer (UV-27F) used in both genotyping and illumina sequencing. LuMiCol isolates sequences were trimmed to match the length of Illumina reads and dereplicated using vsearch. This set of sequences were merged with the dereplicated Illumina reads resulting in merged uniques. The merged fasta file was used for generation of centroids (98% identity), chimera detection and mapped to the whole dataset.

#### Prevalence and absolute abundance analysis

Prevalence was informed by the incidence of each OTU across all samples in the cohort. This was calculated by using the function *amp_core* from ampvis2 v 2.3.2. Output table consists of serial number, OTU number, Frequency (overall), frequency at 1% relative abundance (freq_A), Abundance (mean relative) followed by Taxonomy. Absolute abundance was calculated by using the *phyloseq* object with relative abundance OTU table and multiplying each OTU in each sample by the 16S rRNA gene copies detected per millilitre of BALF sample, quantified using quantitative real-time PCR, using the function:

For each OTU in each sample:

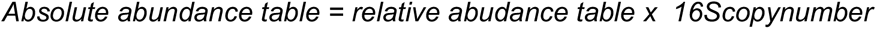

#### Alpha diversity analysis

Alpha diversity indices was obtained from Rényi diversity and corresponding Hill numbers using the function ’renyi’ from *vegan* package in R. The Hill numbers, H_0_ (Number of species), H_1_ (exponent of Shannon diversity), H_2_ (Inverse Simpson) and Hill_∞_ (Berger-Parker index i.e. 1/max *p_i_* (inverse of diversity of order infinity). Proportion of dominant OTUs (max *p_i_*) was calculated by 1/ Hill_∞_ (maximum proportion of species *i*).

#### Beta diversity analysis

Beta diversity was calculated by applying the ‘distance’ function on the phyloseq object. Presence/absence of OTUs was calculated by using Sørenson’s index, which was interchangeably used with the term Bray Curtis distance (binary = TRUE, calls for function ‘vegdist’ from vegan package). For species abundances, Morista-Horn distance measure (calls for function ‘vegdist’ from vegan package and uses the distance measure ‘horn’) was used. Statistical analysis of beta diversity was peformed by PERMANOVA with adonis function in vegan package in R and multiple comparison was performed by using the wrapper function *pairwise.adonis.* We used 10000 permutations as standard for all our comparisons.

#### Enrichment analysis of OTUs

Similar to prevalence analysis across the entire cohort, enrichment analysis of OTUs was repeated for individual PAMs. The *amp_vis* object was split into PAM groups, and the incidence percentages were then calculated by using the function *amp_core* in ampvis2 package. Prevalence of each OTU in individual PAMs was compared to the entire cohort and 30 most prevalent and/or abundant microbiota members were plotted as described in **Table S2**.

For enrichment analysis of OTU abundances, each PAM was compared to a file containing absolute abundances of the other 3 PAMs. Statistical analysis was performed by ART-ANOVA, with a two factorial design (group= single PAM vs other 3 PAMs, variable = OTU IDs). Marginal means were calculated by using the *emmeans* R package. Pairwise differences were calculated followed by Benjamini-Hochberg multiple testing for False Disovery Rate (FDR). Plotting was limited to the 30 most prevalent and/or abundant microbiota members as described in **Table S2**.

### Quantitative analysis of host gene expression and anellovirus load in BALF

#### BALF cellular RNA extraction and real-time quantitative PCR for gene expression analysis

BALF cell lysates were transferred into a QIAshredder column (Qiagen) for homogenization and total RNA was extracted using RNeasy Mini Kit (Qiagen) according to the manufacturer’s instructions. RNA concentration was determined using a Nanodrop ND-1000 spectrophotometer (Thermo Fisher Scientific, Waltham, MA, USA) and reverse transcription was performed using iScript cDNA Synthesis Kit (Bio-Rad, Hercules, CA, USA). Characterization of BAL fluid cell gene expression profiles was based on multiplex real-time PCR analysis using custom oligonucleotide primers and probes (Microsynth, Balgach, Switzerland) for a set of 31 genes (**Dataset S7**). We used guanine nucleotide-binding protein, beta polypeptide 2-like 1 (GNB2L1) gene as a reference gene, given its high expression stability in BALF cells in both health and disease(Ishii et al., 2006). Amplification was carried out using iQ Multiplex Powermix Master Mix and a CFX96 Real-Time detection system.

For radar chart visualization, the samples were sorted according to their association with one of the four pneumotypes, and the median expression values for each gene were determined within each group. For each gene, the highest median was then arbitrarily set to 1 and plotted as the maximum value in the corresponding chart. The median values obtained within the other groups were normalized accordingly.

#### Quantification of anellovirus load

Based on the tropism of *Anelloviridae* for hematopoietic cells, we quantified the load of this virus family starting from the DNA extracted from total BAL fluid cellular pellet. Absolute quantification of pan-Anelloviridae, *Alpha*-, *Beta*- and *Gammatorquevirus* (**See Dataset S7**) was performed using the CFX96 Real-Time detection system (Bio-Rad) based upon values obtained with a set of purified amplicons used as standards.

### Machine learning and statistical modelling

#### Unsupervised learning for Pneumotype discovery

Pneumotypes were obtained by running k-medoid-based unsupervised machine learning using Bray-Curtis dissimilarity matrix (binary = FALSE) using Genocrunch (See Section - Statistical analysis and softwares. The program utilizes the ‘pamk’ function of the R package ‘fpc’ version 2.1.10 to cluster samples while optimizing the number of clusters based on the average silouhette width (Reynolds et al., 2006; Schubert and Rousseeuw, 2019).

#### Random Forest classification and regression based machine learning

Random forest analysis was performed using median normalized expression of 31 genes in **Figure 4A** from all 234 samples, as predictors. For classification based analysis, the pneumotypes were used as responses. For regression based analysis, bacterial and total anellovirus copy numbers were used as responses. These models were optimized for best accuracy and sensitivity using different combinations of sub-sampling (mtry) and number of decision trees (ntrees) constructed at each step. In each analysis, random forest cross-validates results 10 times by creating random shuffled copies of the data. After each analysis, random forest provides results in terms error rate (out-of-box error) and matrix for the predictions for classifications (**See Table S5**) and percentage variance explained for regressions (**See Figure 4I and 5F**). Importance of predictors was calculated by Boruta, which creates random shuffled copies of the data of all features (shadow features: minimum, mean and maximum). At every iteration, it checks whether the real feature has a higher importance than the best of its shadow features (i.e. whether the feature has a higher Z score than the maximum Z score of its shadow features) and constantly removes features which are deemed highly unimportant. Finally, it assigns predictors with an Importance score and categorizes as Confirmed, Tentative or Rejected.

#### Correlations of gene expression with predicted features by random forest regression

Gene predictors from random forest analysis with importance scores more than 10 were further fitted into additive linear model with either bacterial or viral copy numbers i.e. lm(copy number ∼ gene A + gene B). The best models were selected ’stepAIC’ function from *MASS* package in R, which performs a stepwise model selection by AIC (Akaike Information Criteria).

### Clinical measurements and definitions

Determination of the cell differential in the BALF, B-cell count in peripheral blood by mass cytometry, and bacterial culture for diagnostic purposes were performed according to in-house routine clinical procedures.

#### Definition of acute bacterial infection

Acute bacterial infection was defined as positive BALF culture with dedicated antibiotic treatment, associated with clinical signs and symptoms such as a decrease in FEV1, new or progressive infiltrate on standard chest radiography or CT-scan, fever, positive pulmonary auscultation, cough, dyspnea, hemoptysis, pleuritic pain, purulent sputum.

In contrast, a BALF culture positive for a pathogen, but not associated with the administration of antibiotic therapy and without clinical signs and/or symptoms, was considered as a bacterial colonization and not as an acute bacterial infection.

#### Definition of Chronic Lung Allograft dysfunction (CLAD)

CLAD was defined as a loss of more than 20% of the expiratory volume in 1 second (FEV1) of the mean of the two best values (i.e. the baseline FEV1) since transplantation, without other obvious cause and without reversibility, in accordance with the diagnostic criteria specified by the Pulmonary Council of the International Society for Heart and Lung Transplantation (Verleden et al., 2020)

## Supporting information

Supplementary Datasets (ZIP file)

## Acknowledgements

We thank the clinicians of the Pulmonology Division of CHUV who performed the bronchoscopies, the clinical database team of the Thoracic Surgery Division of CHUV (Prof Thorsten Krueger, Audrey Roth RN and Fébronie Maillefer RN) for contributing in the completion of the clinical dataset, Julie Pernot for technical assistance, and the transplant patients for allowing their clinical data and BALF samples to be used for clinical research.

## Funding

During this project, SD received the European Union Marie Sklodowska-Curie Individual Fellowship with the project HUMANITY (Grant Agreement no. 800301) and is hosted by PE. PE & BM received the Interdisciplinary grant (from the Faculty of Biology and Medicine, University of Lausanne, Grant No. 26075716). PE received the ERC StG (MicroBeeOme, Grant No. 714804) and SNSF project (Grant no. 31003A_179487). AK received grants from the Swiss Lung Association, Bern (Research Fund No 2018-16), the University of Lausanne / Centre Hospitalier Universitaire Vaudois (Grant Pépinière) and GSK educational grant. EB was supported by “Fondation Professeur Placide Nicod”.

## Author Contributions

Conceptualization, S.D., E.B., D.-A.W., J.-D.A., B.J.M., P.E. and L.P.N.; Data Curation, S.D., E.B., A.K., J.-D.A. and V.T.; Formal Analysis, S.D., E.B. and D.-A.W.; Funding Acquisition, S.D., A.K., C.V.G., B.J.M., P.E. and L.P.N.; Investigation, S.D., E.B., D.-A.W., V.T., J.-D.A., M.-F.D., L.M. and C.P.; Methodology, S.D., E.B., D.-A.W., V.T. and G.B.-R.; Project Administration, E.B., A.K., B.J.M., P.E. and L.P.N.; Resources, A.K., C.P., A.R., C.V.G., B.J.M., P.E. and L.P.N.; Software, S.D. and G.B.-R.; Supervision, E.B., A.K., G.B.-R., C.V.G., B.J.M., P.E. and L.P.N.; Validation, S.D., E.B., D.-A.W., V.T., J.-D.A. and P.E.; Visualization, S.D., E.B. and D.-A.W.; Writing - Original Draft, S.D., E.B., and P.E.; Writing - Review & Editing, S.D., E.B., A.K., D.-A.W., V.T., G.B.-R., J.-D.A., M.-F.D., L.M., C.P., A.R., C.V.G., B.J.M., P.E. and L.P.N.

## Figures

**Figure S1.**
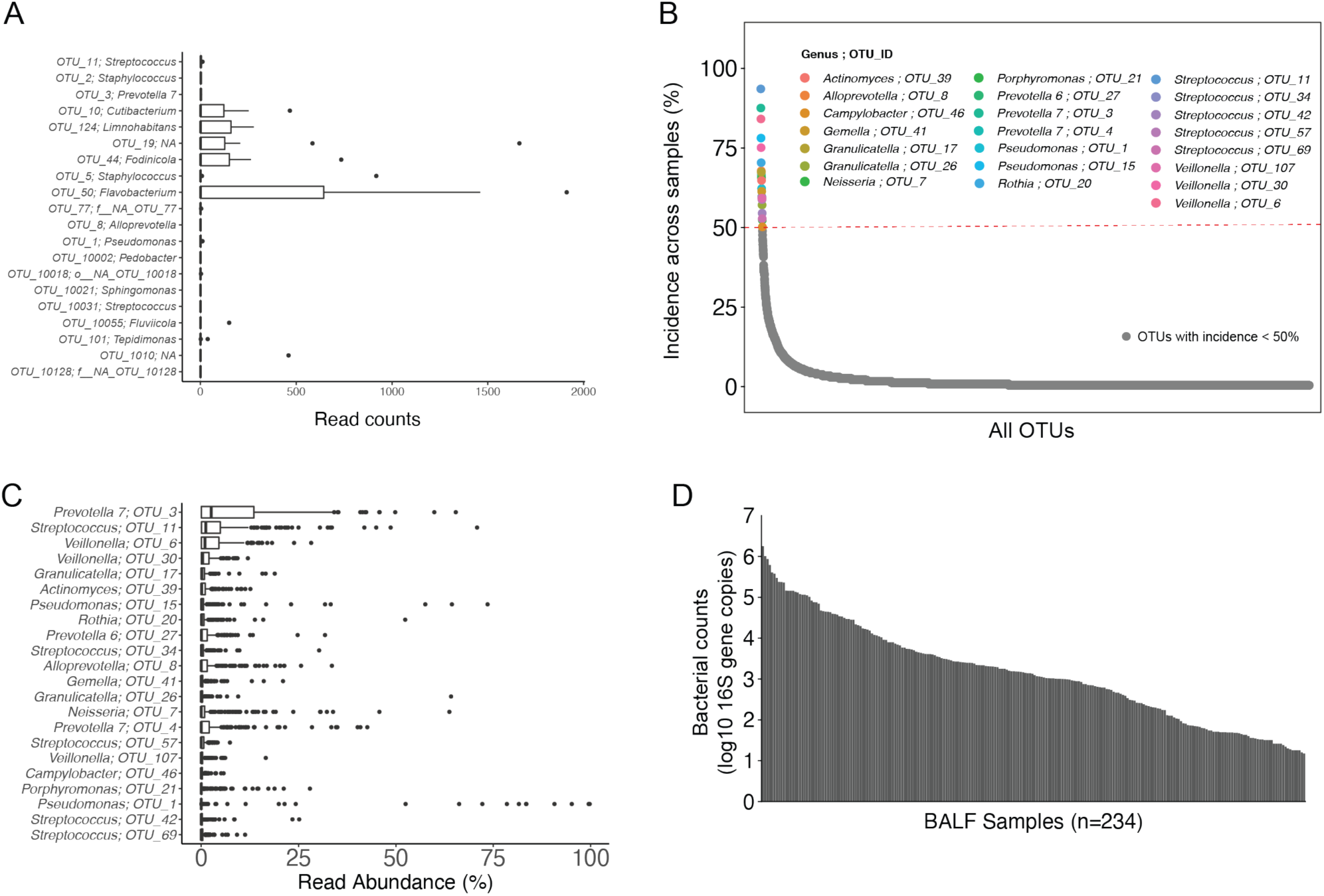
Relative abundance and prevalence analyses of OTUs detected in the negative controls as well as of the most abundant OTUs across all 234 BALF samples. **(A)** Box plot showing number of reads per BALF samples contributed by ambiguous OTUs also detected in negative control samples, which included Bronchoscope pre-wash, DNA extraction reagents and no-template PCR reaction. **(B)** Incidence plot showing the frequencies of OTUs across all BALF samples (%, y-axis) in our cohort (x-axis). Colored points show genera and OTU IDs of the taxa present in ≥50% of BALF samples (red dotted line), while grey points show OTUs with incidences ≤ 50%. **(C)** Box plot showing relative abundances (%) of the most abundant OTUs (denoted by genera and OTU IDs, median abundance) across all BALF samples.

**Figure S2.**
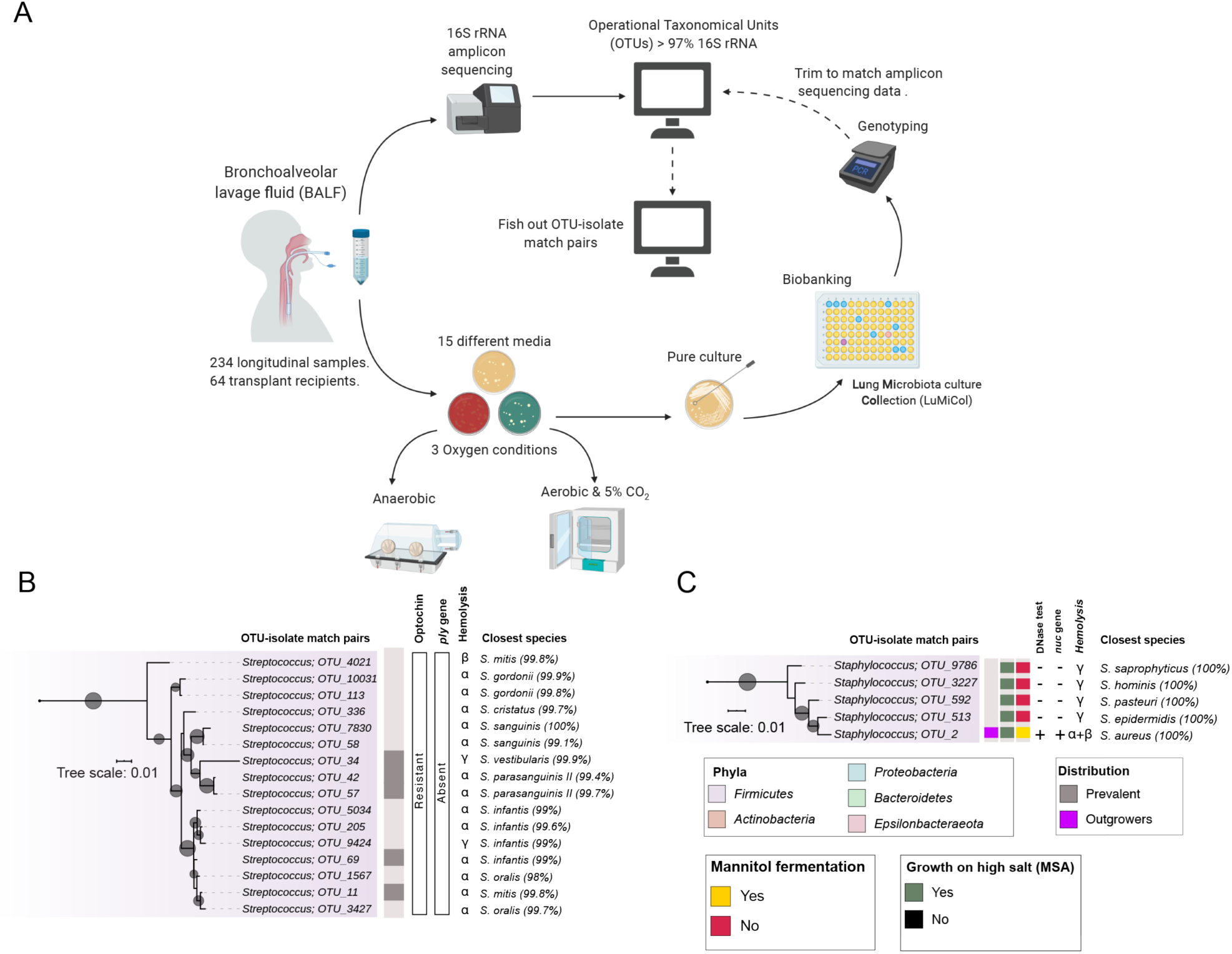
Workflow of our combined approach of BALF amplicon sequencing and culturomics to deduce the microbial ecology of deep lung microbiota. **(A)** Amplicon sequencing of the 16S rRNA gene was carried out for 234 bronchoalveolar lavage fluid (BALF) samples from 64 recipients post-lung transplant. The resulting reads were clustered into operational taxonomic units (OTUs) to determine the community composition of each sample. For a subset of the samples, bacteria were cultured on 15 different media and 3 oxygen conditions. Single colonies were picked, genotyped, and arrayed into a bacterial strain collection referred to as LuMiCol. 16S rRNA gene sequences of these isolates were included into the community analysis based on the culture-independent 16S rRNA gene amplicon sequences to identify which isolate belongs to which OTU (OTU-isolate matching pairs). (B and C) Phenotypic tests to differentiate bacteria species. Taxa are denoted by genera and OTU IDs and phyla are shown with colored highlights and were also assigned if they are prevalent (grey rectangle) or opportunists (magenta rectangle) of the lower respiratory community. Streptococci were confirmed by their characteristic hemolysis (C). Viridans Streptococci were differentiated from Pneumococcus by Optochin resistance test and the presence of *ply* gene encoding Pneumolysin toxin. *Staphylococcus aureus* was differentiated from other Staphylococci (D) by its ability to grow in high salt concentration and fermentation of Mannitol (Mannitol Salt Agar), characteristic hemolysis, presence of *nuc* gene encoding for Staphylococcal Thermonuclease and extracellular DNase activity assay.

**Figure S3.**
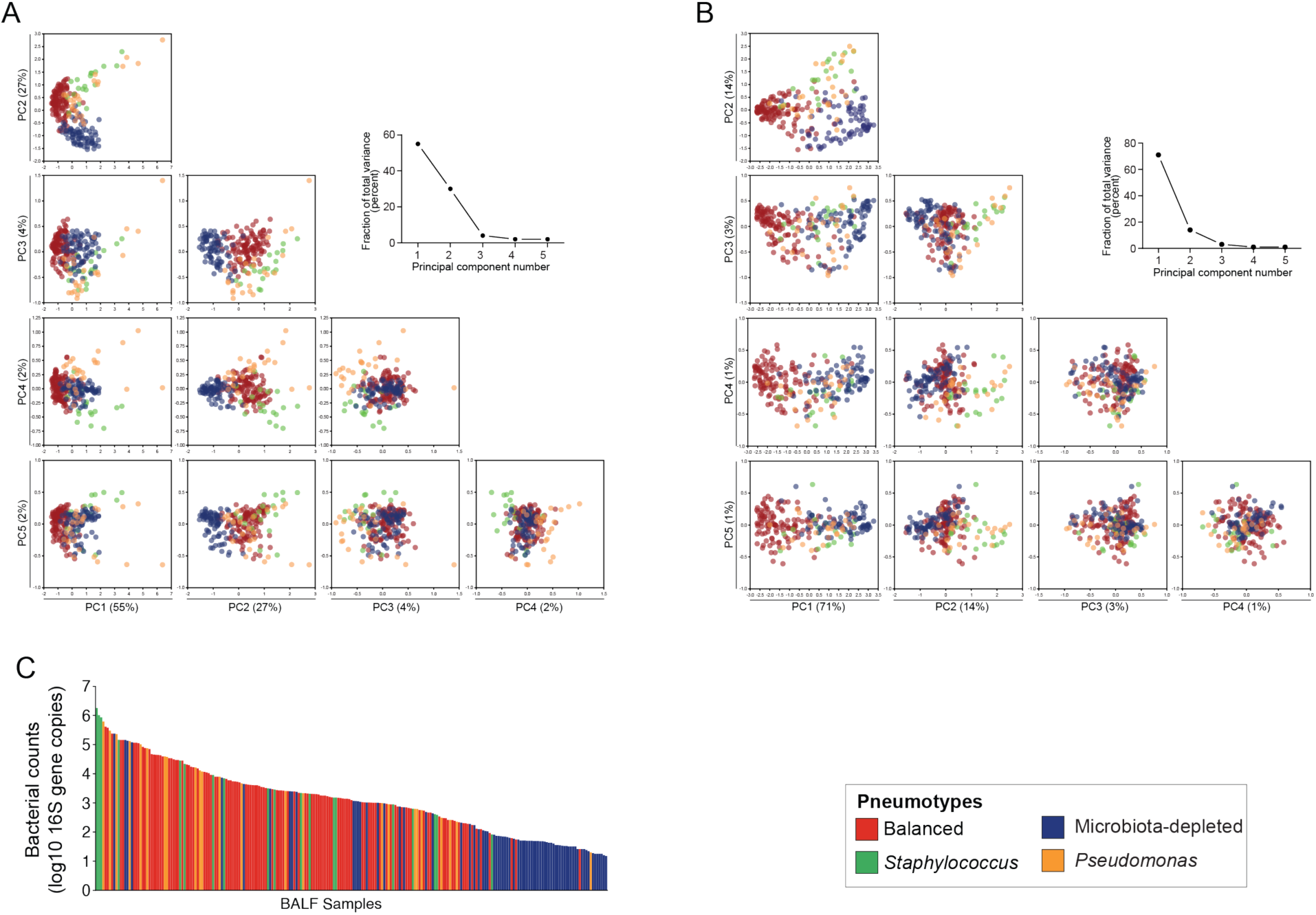
Principal component analysis based on the bacterial community compositions and bacterial loads of the 234 BALF samples. **(A and B)** Clustering of samples based on genus level (A) and OTU level (B) along principal components 1 to 5. Samples are colored according to PAM designation. Line graph shows total variance explained per principal component. **(C)** Bacterial count as determined by qPCR on the 16S rRNA gene. Samples are sorted according to load and colors correspond to PAM designation. PAM1, PAM2, PAM3, and PAM4 correspond to red, green, blue, and orange colors, respectively.

**Figure S4.**
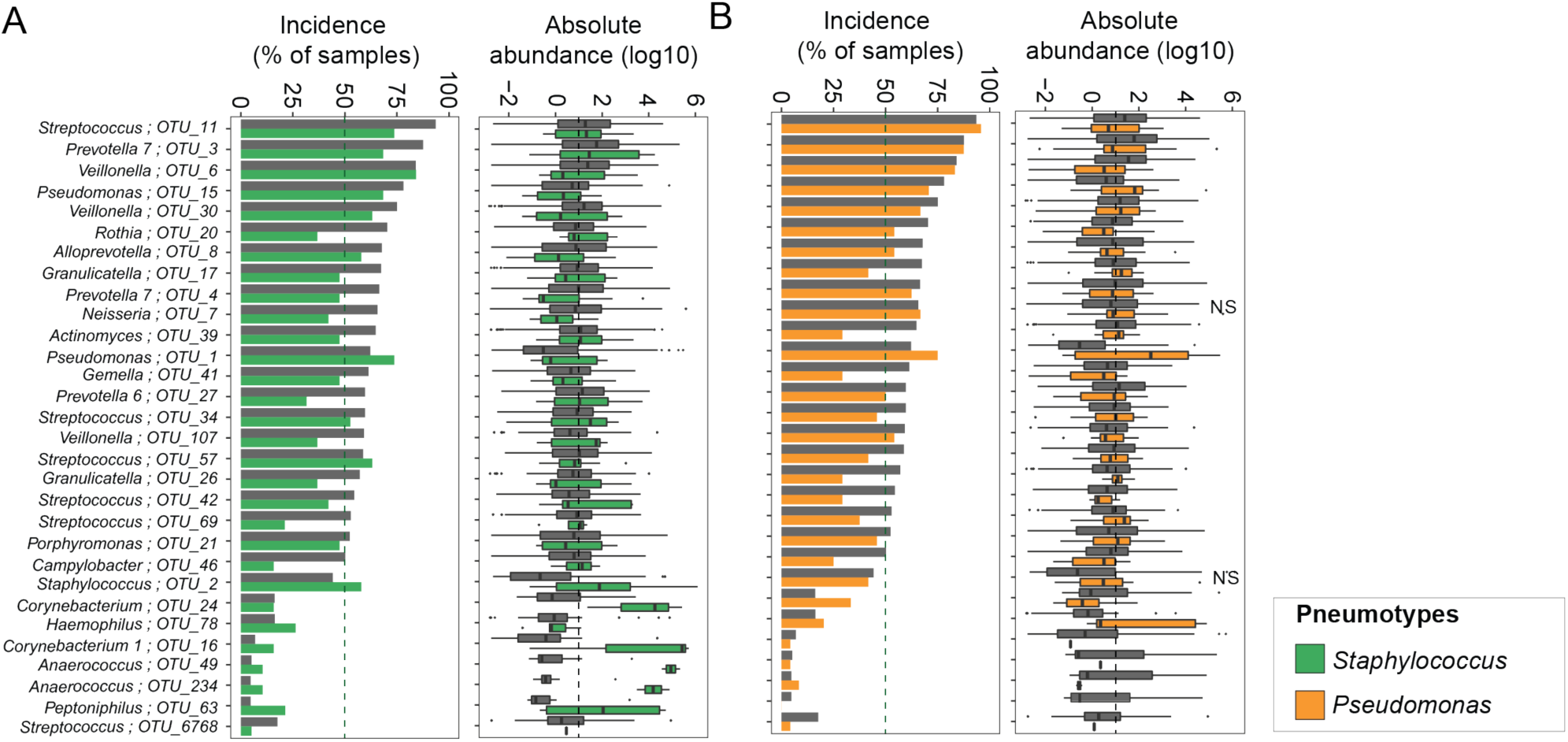
Prevalence (i.e % of samples present) and absolute abundance across all samples of the 30 most dominant bacterial community members (i.e. OTUs) in PAM2. **(A)** and PAM4 (B) as compared to the other 3 PAMs. Green and orange graphs correspond to values for PAM2 and PAM4, respectively, while grey graphs correspond to values for the entire dataset. Incidence of 50% is indicated by the green dotted line. Enrichment analysis was performed on both incidences and abundances of specific bacteria in individual PAMs compared to the other 3 PAMs. Differential abundances were analyzed by ART-ANOVA followed by Benjamini-Hochberg multiple testing for False Discovery Rate (FDR) (only NS = Not significant are shown).

**Figure S5.**
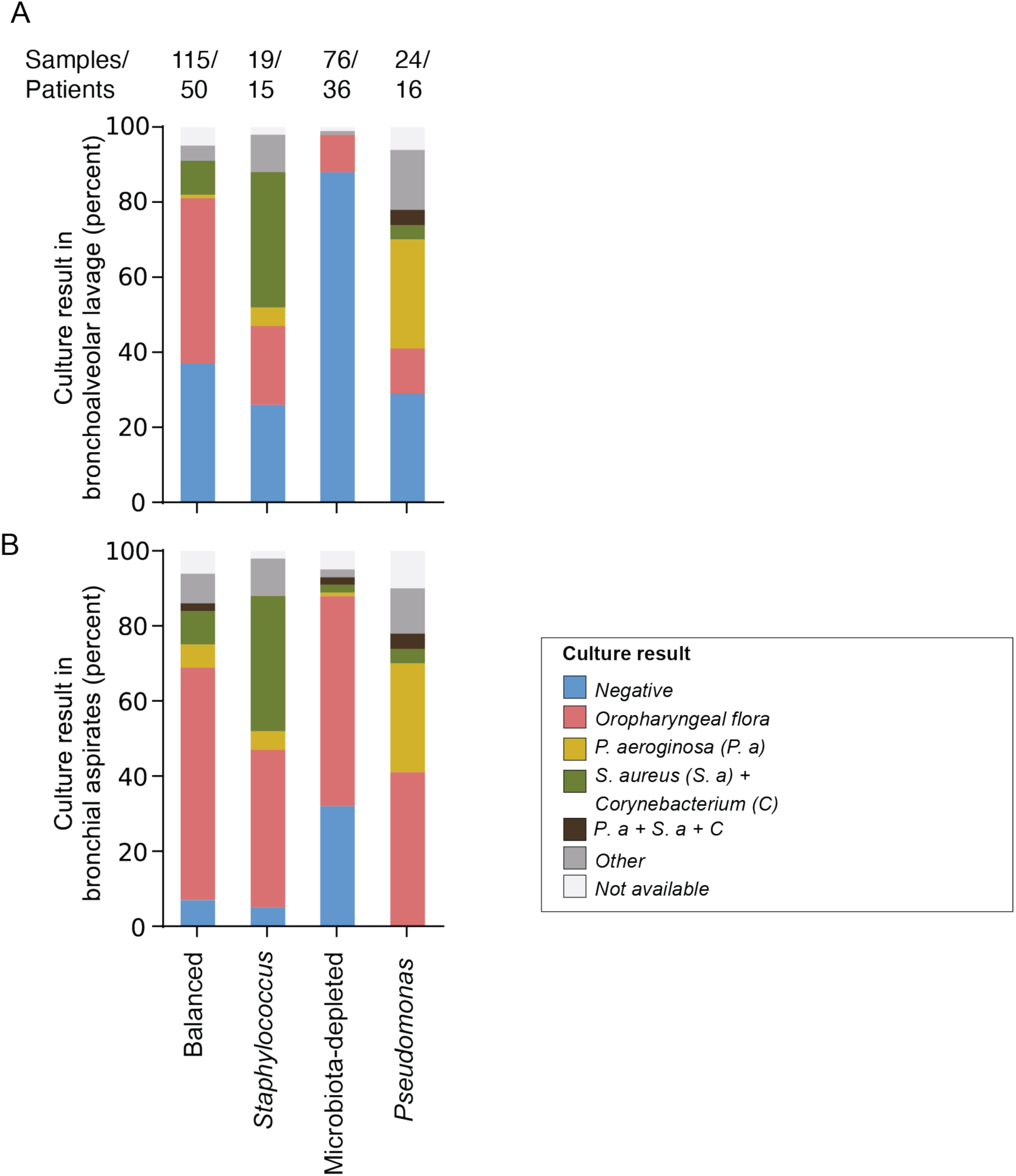
Comparison of culturing results from BALF and Bronchial aspirate (BAs) samples belonging to different pneumotypes (as based on BALF community analysis) confirms the presence of distinct microbial communities. (A and B) Stacked bar plots shows relative percentage of specific taxonomic groups (colored bars) isolated from paired Bronchial aspirates (BA) and Bronchoalveolar lavage fluid (BALF) samples, plotted according to the Pneumotype designation of each sample pair.

**Figure S6.**
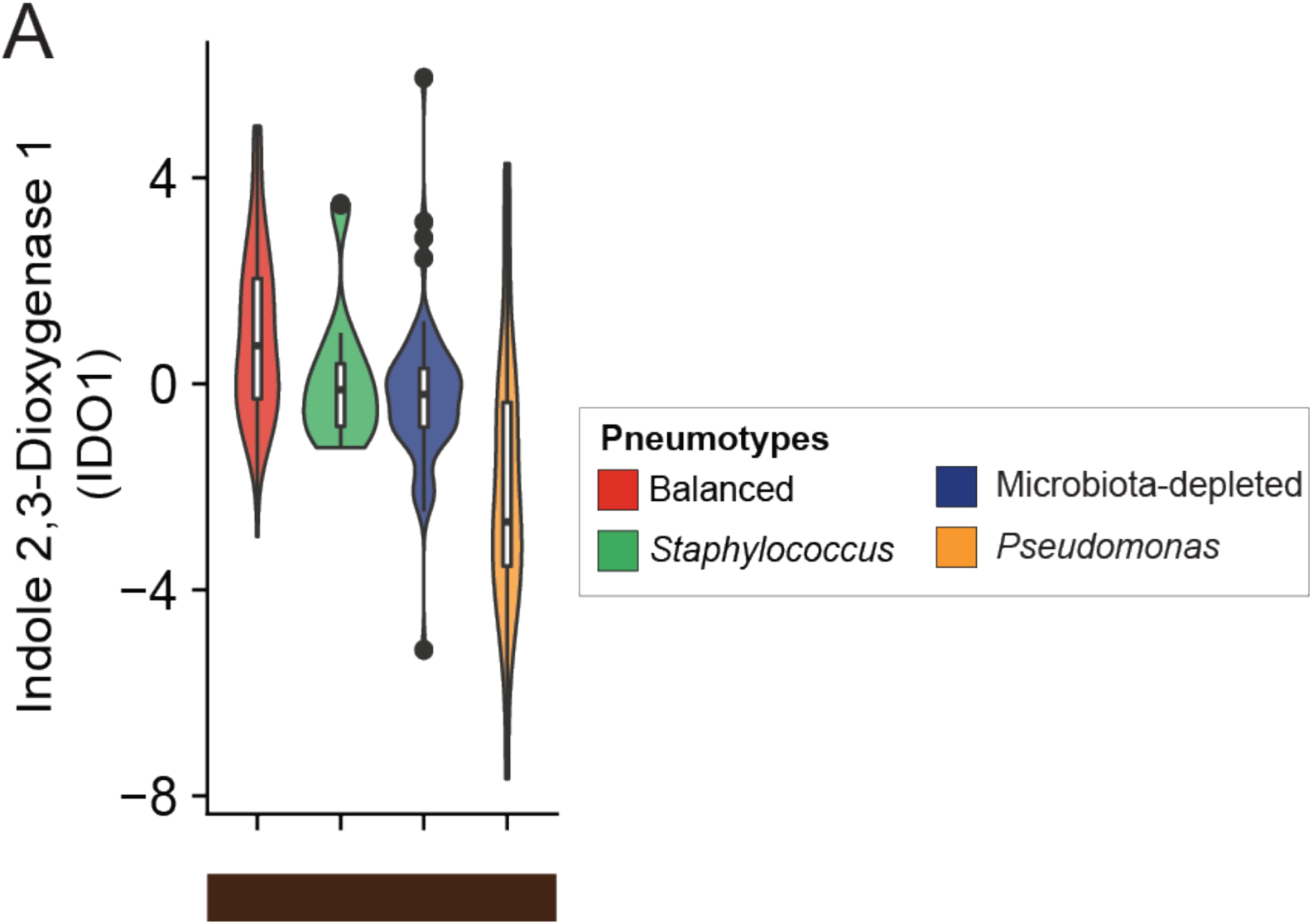
Gene expression differences of peripheral immune tolerance gene IDO1 across the four pneumotypes. Violin plots showing median normalized expression of IDO1 (log2 fold) relative to reference gene (GNB2L1, Methods) across the four pneumotypes (plot colors). The functional category is shown at the bottom of the plot according to the color scheme used in Figure 4A. For statistical analysis between groups, data normality was checked by Levene’s test followed by either one-way ANOVA followed by Tukey’s post hoc test or Kruskal-Wallis test followed by Dunn’s post hoc test.

**Figure S7.**
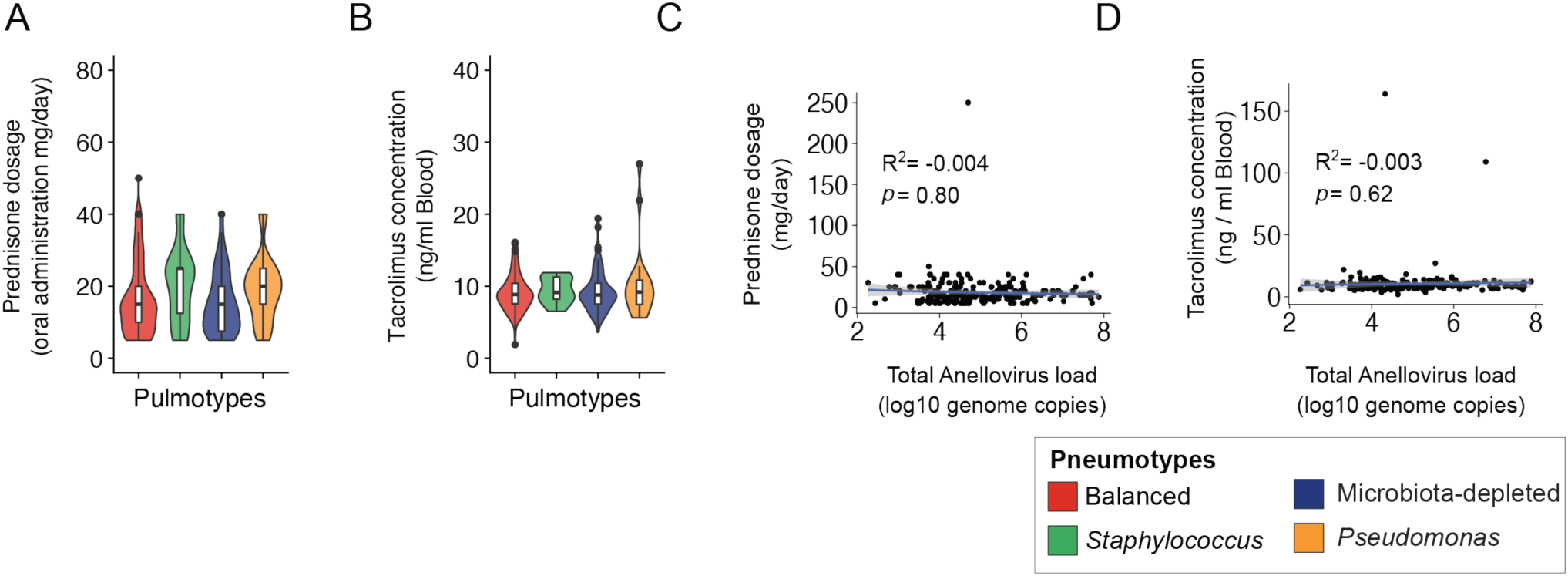
Association of Immunosuppresant drugs with pneumotypes and anellovirus load in BALF. (**A and B**) Violin plots showing levels of immunosuppresants: Prednisone dosage (mg/day) and Tacrolimus concentration in blood (ng/ml) (y-axis) in BALF samples from four pneumotypes (plot colors and x-axis). (**C and D**) Scatter plot showing correlation between levels of immunosuppresants: Prednisone dosage (mg/day) and Tacrolimus concentration in blood (ng/ml) (y-axis) and Total anellovirus burden (log 10 genome copies per ml BALF per sample (x-axis). R^2^ indicates the proportion of explained variability with significance given by *p* value.

**Figure S8.**
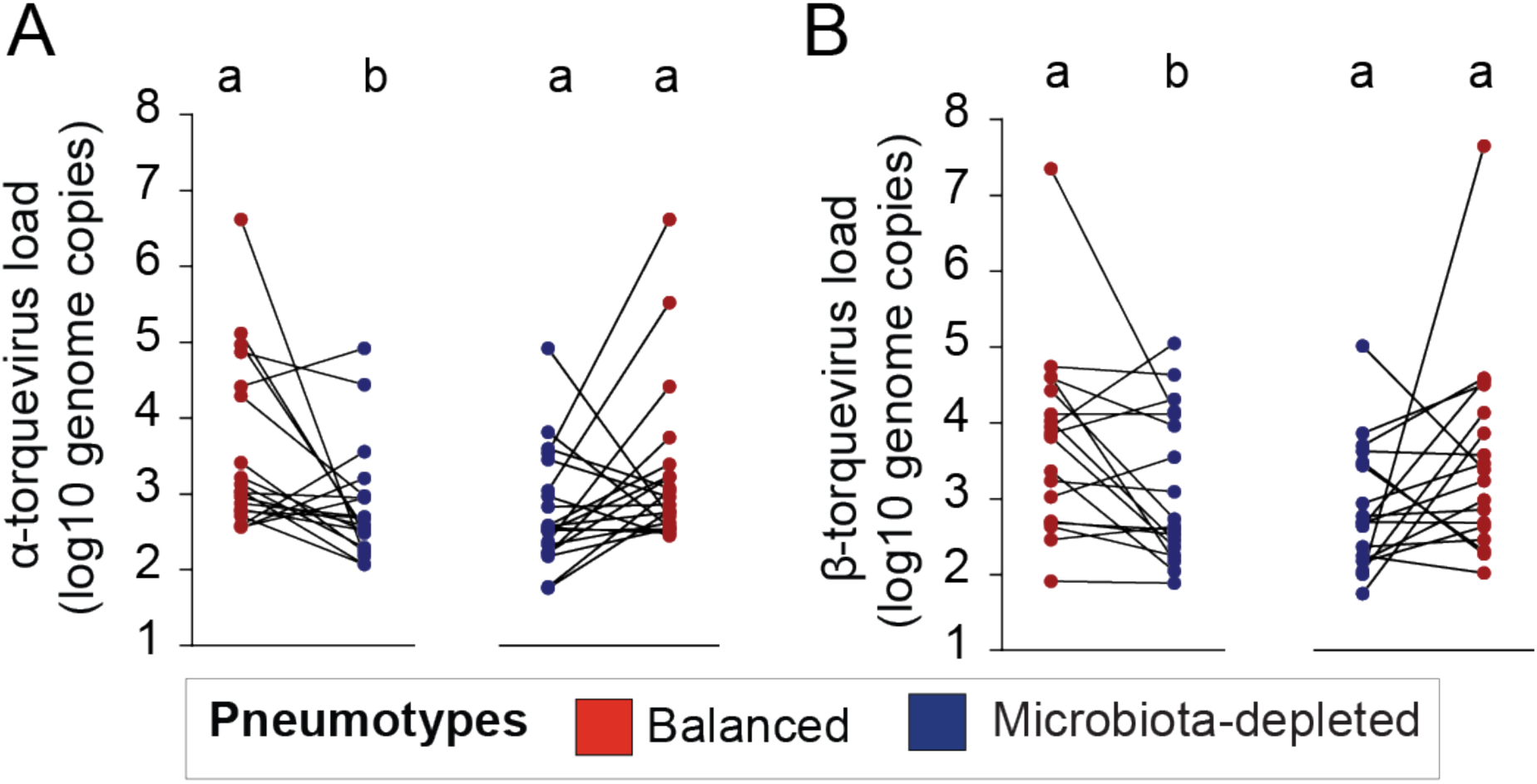
Burden of major anellovirus genera in BALF differ between pneumotypes. Intra-individual pairwise analysis of ⍺- and β-torquevirus loads (log 10 genome copies per ml BALF) for the samples transitioning from Pneumotype_Balanced_ (Red) to Pneumotype_MD_ (Blue) and from Pneumotype_MD_ to PneumotypeBalanced. Statistical analysis was performed by paired Wilcoxon Signed Rank sum test. Different letters denote signficant differences between the two pneumotypes.

**Figure S9.**
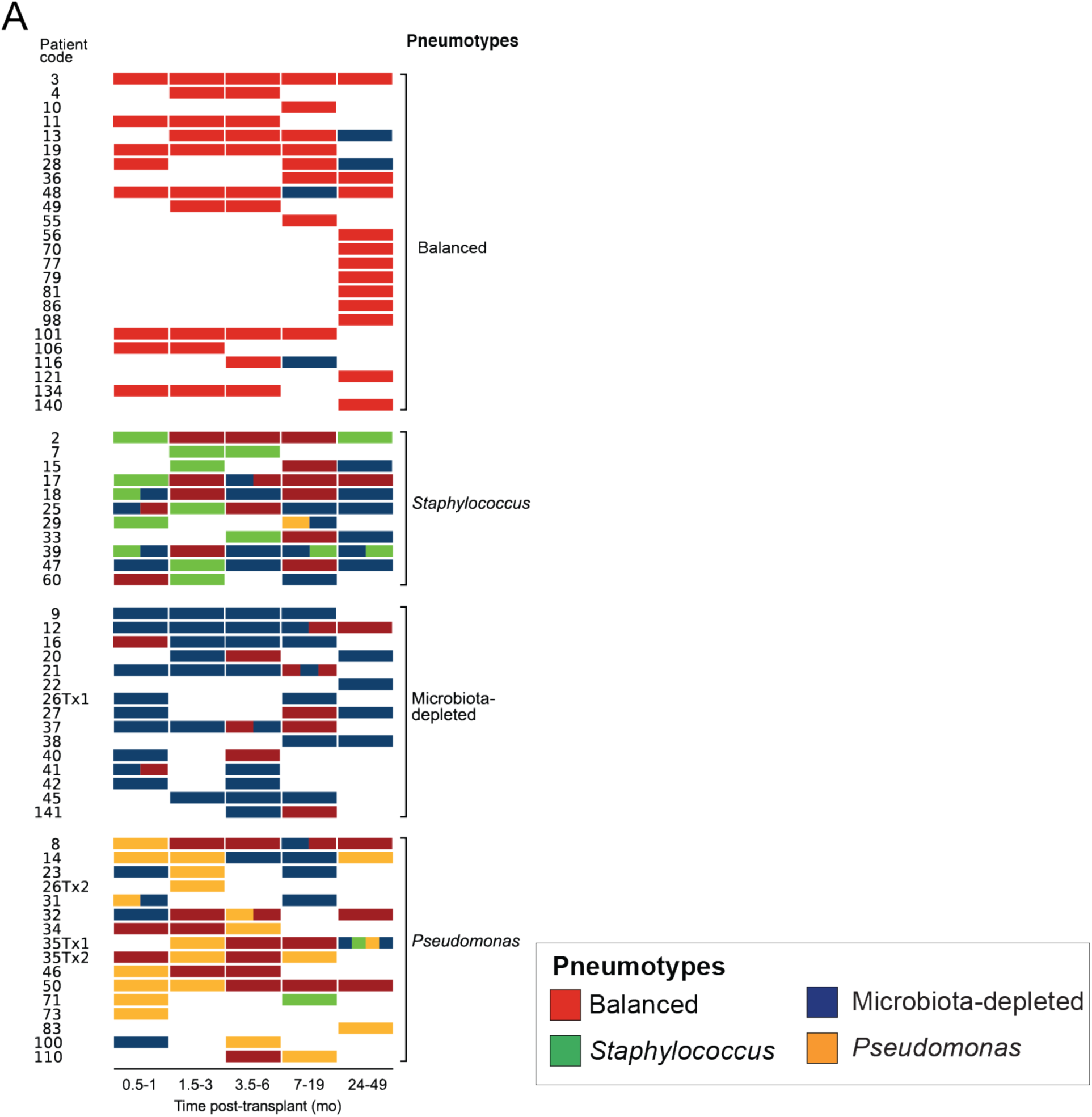
Longitudinal analysis of lung microbiota post-transplant to investigate pneumotype transitions. **(A)** Longitudinal BALF sampling from patients with given IDs and integrated pneumotype information across time (months post-transplant).

## Supplementary Tables and Datasets

Larger Tables are provided as Datasets are available online on the cloud drive. https://drive.switch.ch/index.php/s/hch0EoA5QyjBPR8

**Dataset S1**: 16S rRNA amplicon sequencing full table with sample-wise information OTUs relative abundance and taxonomy, without negative controls.

**Dataset S2**: Negative controls sequencing full table with sample-wise information OTUs abundance and taxonomy.

**Dataset S3**: Detailed table showing frequency of all OTUs detected across the 234 BALF samples in terms of their abundance and prevalence across samples.

**Dataset S4**: Overview of the different combinations of culture conditions used in our culturomics approach and source or references for each media.

**Dataset S5**: Lung microbiota culture collection (LuMiCol) isolate list with detailed information about sample number, culture conditions, taxonomy. This is an uncurated list containing approx. 300 isolates including 215 isolates used for analysis.

**Dataset S6**: OTU-isolate match summary including the isolates that match OTUs in 16S rRNA and information about number of representative isolates in LuMiCol, with at least one representative isolate name, prevalence, and oxygen and media preferences

**Dataset S7:** Primer sequences and qPCR specifications for each primer pair used to analyze host gene expression, anelloviruses in BALF and genotyping of bacterial species.

**Dataset S8:** Detailed metadata table associated with all patients and samples.

---

**Table S1:** BALF sample characteristics and patient demographics.

|  |  |  |
| --- | --- | --- |
| **Patients/Samples** | Total (n) | 64/234 |
|  | Male / Female | 25 (39.1%)/39 (60.9%) |
|  | Age at transplant (yr) | 54 (36-60) |
|  | Sampling time point (months post-transplant) | 6 (2-12) |
| **Type of transplant** | Bilateral lung | 61 (95.3%) |
|  | Single lung | 3 (4.7%) |
| **Pre-transplant diagnosis** | Interstitial lung disease | 17 (26.6%) |
|  | Cystic fibrosis | 14 (21.9%) |
|  | Chronic obstructive pulmonary disease | 17 (26.6%) |
|  | Pulmonary hypertension | 4 (6.3%) |
|  | Alpha-1 antitrypsin deficiency | 4 (6.3%) |
|  | Other | 8 (12.5%) |
| **Transbronchial biopsies<sup>a,b</sup>** |  | 224 (95.7%) |
|  | A0 | 182 (77.8%) |
|  | A1 | 21 (9.0%) |
|  | A2 | 5 (2.1%) |
|  | B0 | 148 (63.2%) |
|  | B1 | 39 (16.7%) |
|  | B2 | 2 (0.9%) |
| **Immunosuppression<sup>a</sup>** | Tacrolimus | 231 (98.7%) |
|  | Cyclosporin | 2 (0.9%) |
|  | Everolimus | 1 (0.4%) |
| **Antibiotics<sup>a,c</sup>** | PCP prophylaxis (TMP/SMX/Atovaquone) | 218 (93.1%) |
|  | Inhaled (colistin, ambisome, tobramycin) | 24 (10.3%) |
|  | Oral or IV route (miscellaneous, including azithromycin) | 71 (30.3%) |
| **Clinical infection<sup>a,d</sup>** |  | 33 (14.1%) |

Abbreviations: IV=intravenous; PCP=pneumocystis, pneumonia; TMP/SMX=trimethoprim/sulfamethoxazole.
Data presented as n (% of patients or samples) or median (interquartile range)
<sup>a</sup> at sampling
<sup>b</sup> grading of pulmonary allograft rejection according to guidelines of the International Society for Heart and Lung Transplantation
<sup>c</sup> antibacterial and antifungal antibiotics
<sup>d</sup> BALF positive culture requiring treatment

**Table S2:** Incidences and abundances of 30 most prevalent and/or abundant microbiota members of human lower respiratory tract

| OTU_ID | Genera | I (%) | Ai (%) | Mean abundance (relative %) | Cumulative abundance (absolute, %) | Present in LuMiCol |
| --- | --- | --- | --- | --- | --- | --- |
| OTU_11 | *Streptococcus* | 93.6 | 54.1 | 5.1 | 2.0 | Yes |
| OTU_3 | *Prevotella 7* | 87.6 | 57.9 | 10.3 | 9.1 | Yes |
| OTU_6 | *Veillonella* | 84.1 | 49.8 | 3.7 | 1.6 | Yes |
| OTU_15 | *Pseudomonas* | 78.1 | 19.3 | 2.5 | 1.1 | No |
| OTU_30 | *Veillonella* | 75.1 | 37.3 | 1.8 | 0.9 | Yes |
| OTU_20 | *Rothia* | 70.4 | 21.0 | 1.5 | 0.3 | Yes |
| OTU_8 | *Alloprevotella* | 67.8 | 27.5 | 3.3 | 1.0 | No |
| OTU_17 | *Granulicatella* | 67.4 | 21.5 | 1.3 | 0.6 | Yes |
| OTU_4 | *Prevotella 7* | 66.5 | 30.5 | 4.4 | 1.7 | No |
| OTU_7 | *Neisseria* | 65.7 | 24.5 | 3.8 | 6.0 | Yes |
| OTU_39 | *Actinomyces* | 64.8 | 25.3 | 1.5 | 1.1 | Yes |
| OTU_1 | *Pseudomonas* | 62.2 | 10.3 | 7.0 | 7.5 | Yes |
| OTU_41 | *Gemella* | 61.4 | 14.2 | 1.0 | 0.1 | Yes |
| OTU_27 | *Prevotella 6* | 59.7 | 31.8 | 2.3 | 0.7 | Yes |
| OTU_34 | *Streptococcus* | 59.7 | 18.9 | 1.2 | 0.1 | Yes |
| OTU_107 | *Veillonella* | 59.2 | 10.3 | 0.8 | 0.3 | Yes |
| OTU_57 | *Streptococcus* | 58.8 | 19.3 | 0.9 | 0.4 | Yes |
| OTU_26 | *Granulicatella* | 57.1 | 9.9 | 1.1 | 0.3 | No |
| OTU_42 | *Streptococcus* | 54.5 | 11.2 | 1.1 | 0.2 | Yes |
| OTU_69 | *Streptococcus* | 52.8 | 12.9 | 0.9 | 0.1 | Yes |
| OTU_21 | *Porphyromonas* | 52.4 | 16.3 | 2.3 | 1.4 | No |
| OTU_46 | *Campylobacter* | 50.2 | 10.3 | 0.6 | 0.3 | No |
| OTU_2 | *Staphylococcus* | 44.2 | 6.0 | 5.6 | 22.0 | Yes |
| OTU_6768 | *Streptococcus* | 17.6 | 1.3 | 0.8 | 1.1 | No |
| OTU_24 | *Corynebacterium*<br>1 | 16.3 | 2.1 | 5.4 | 3.3 | No |
| OTU_78 | *Haemophilus* | 16.3 | 1.3 | 3.0 | 1.3 | No |
| OTU_16 | *Corynebacterium*<br>1 | 6.9 | 1.3 | 4.4 | 9.3 | No |
| OTU_49 | *Anaerococcus* | 5.2 | 1.3 | 4.4 | 2.9 | No |
| OTU_234 | *Anaerococcus* | 4.7 | 1.2 | 1.02 | 0.9 | No |
| OTU_63 | *Peptoniphilus* | 4.7 | 1.2 | 2.4 | 0.9 | No |

I Overall incidence of OTUs (% of samples).
Ai Incidence of OTUs at 1% relative abundance.

**Table S3:** Beta diversity summary and statistics of four Partition around medoids (PAMs) formed by samples from the human lower respiratory tract

| **Pairs** | ***p* value** | ***adj. p* value** | **Significance** | **Beta diversity measure** |
| --- | --- | --- | --- | --- |
| PAM1 vs PAM2 | 0.001 | 0.0012 | ** | Sorenson Index |
| PAM1 vs PAM3 | 0.001 | 0.0012 | ** | Sorenson Index |
| PAM1 vs PAM4 | 0.001 | 0.0012 | ** | Sorenson Index |
| PAM2 vs PAM3 | 0.001 | 0.0012 | ** | Sorenson Index |
| PAM2 vs PAM4 | 0.067 | 0.067 | ns | Sorenson Index |
| PAM3 vs PAM4 | 0.001 | 0.0012 | ** | Sorenson Index |
| PAM1 vs PAM2 | 0.001 | 0.0012 | ** | Unweighted UniFrac |
| PAM1 vs PAM3 | 0.001 | 0.0012 | ** | Unweighted UniFrac |
| PAM1 vs PAM4 | 0.001 | 0.0012 | ** | Unweighted UniFrac |
| PAM2 vs PAM3 | 0.001 | 0.0012 | ** | Unweighted UniFrac |
| PAM2 vs PAM4 | 0.091 | 0.091 | ns | Unweighted UniFrac |
| PAM3 vs PAM4 | 0.001 | 0.0012 | ** | Unweighted UniFrac |
| PAM1 vs PAM2 | 0.001 | 0.0015 | ** | Weighted UniFrac |
| PAM1 vs PAM3 | 0.001 | 0.0015 | ** | Weighted UniFrac |
| PAM1 vs PAM4 | 0.001 | 0.0015 | ** | Weighted UniFrac |
| PAM2 vs PAM3 | 0.002 | 0.0024 | ** | Weighted UniFrac |
| PAM2 vs PAM4 | 0.021 | 0.021 | * | Weighted UniFrac |
| PAM3 vs PAM4 | 0.001 | 0.0015 | ** | Weighted UniFrac |

**Table S4:** Statistical evaluation of bacteria composition of PAM1

| **Taxonomy** | **Incidence (%)** | **Abundance (Enriched/Reduced)** | **Significance (Abundance)** | **Present in LuMiCol** |
| --- | --- | --- | --- | --- |
| *Streptococcus* ; OTU_11 | 99.1 | Enriched | *** | Yes |
| *Prevotella* 7 ; OTU_3 | 97.4 | Enriched | *** | Yes |
| *Veillonella* ; OTU_6 | 93.9 | Enriched | *** | Yes |
| *Veillonella* ; OTU_30 | 93 | Enriched | NS | Yes |
| *Granulicatella* ; OTU_17 | 93 | Enriched | ** | Yes |
| *Actinomyces* ; OTU_39 | 89.6 | Enriched | NS | Yes |
| *Rothia* ; OTU_20 | 88.7 | Enriched | *** | Yes |
| *Granulicatella* ; OTU_26 | 84.3 | Enriched | *** | No |
| *Gemella* ; OTU_41 | 83.5 | Enriched | *** | Yes |
| *Neisseria* ; OTU_7 | 82.6 | Enriched | *** | Yes |
| *Prevotella* 7 ; OTU_4 | 81.7 | Enriched | *** | No |
| *Prevotella* 6 ; OTU_27 | 81.7 | Enriched | NS | Yes |
| *Streptococcus* ; OTU_57 | 81.7 | Enriched | ** | Yes |
| *Pseudomonas* ; OTU_15 | 80.9 | Reduced | *** | No |
| *Alloprevotella* ; OTU_8 | 79.1 | Enriched | *** | No |
| *Streptococcus* ; OTU_34 | 79.1 | Enriched | *** | Yes |
| *Veillonella* ; OTU_107 | 76.5 | Enriched | *** | Yes |
| *Streptococcus* ; OTU_69 | 76.5 | Enriched | *** | Yes |
| *Campylobacter* ; OTU_46 | 75.7 | Enriched | *** | No |
| *Streptococcus* ; OTU_42 | 73.9 | Enriched | *** | Yes |
| *Porphyromonas* ; OTU_21 | 67 | Enriched | NS | No |
| *Pseudomonas* ; OTU_1 | 59.1 | Reduced | *** | Yes |

**Table S5:** Random forest confusion matrix for prediction of pneumotypes using host immune gene expression.

| **Pneumotypes** | **Balanced** | ***Staphylococcus*** | **Microbiota-depleted** | ***Pseudomonas*** | **Accuracy (%)** |
| --- | --- | --- | --- | --- | --- |
| **Balanced** | 103 | 0 | 8 | 1 | 92% |
| ***Staphylococcus*** | 5 | 0 | 13 | 0 | 0% |
| **Microbiota-depleted** | 13 | 0 | 61 | 1 | 81.4% |
| ***Pseudomonas*** | 5 | 0 | 0 | 19 | 83.4% |

## Notes

https://drive.switch.ch/index.php/s/hch0EoA5QyjBPR8

### Competing Interest Statement

The authors have declared no competing interest.

## References

Abbas, A.A., Diamond, J.M., Chehoud, C., Chang, B., Kotzin, J.J., Young, J.C., Imai, I., Haas, A.R., Cantu, E., Lederer, D.J., et al. (2017). The Perioperative Lung Transplant Virome: Torque Teno Viruses Are Elevated in Donor Lungs and Show Divergent Dynamics in Primary Graft Dysfunction. Am. J. Transplant. 17, 1313–1324.

Andersen, S.K., Kirkegaard, R.H., Karst, S.M., and Albertsen, M. (2018). ampvis2: an R package to analyse and visualise 16S rRNA amplicon data. BioRxiv.

Arend, W.P., Malyak, M., Guthridge, C.J., and Gabay, C. (1998). INTERLEUKIN-1 RECEPTOR ANTAGONIST: Role in Biology. Annu. Rev. Immunol. 16, 27–55.

Arumugam, M., Raes, J., Pelletier, E., Le Paslier, D., Yamada, T., Mende, D.R., Fernandes, G.R., Tap, J., Bruls, T., Batto, J.M., et al. (2011). Enterotypes of the human gut microbiome. Nature.

Atarashi, K., Tanoue, T., Oshima, K., Suda, W., Nagano, Y., Nishikawa, H., Fukuda, S., Saito, T., Narushima, S., Hase, K., et al. (2013). Treg induction by a rationally selected mixture of Clostridia strains from the human microbiota. Nature.

Awasthi, S., Singh, B., Ramani, V., Xie, J., and Kosanke, S. (2019). TLR4-interacting SPA4 peptide improves host defense and alleviates tissue injury in a mouse model of Pseudomonas aeruginosa lung infection. PLoS One.

Beaume, M., V., L., T., K., N., G., O., M., J.-D., A., L., B., L., F., P., G., J., S., et al. (2016). Microbial communities of conducting and respiratory zones of lung-transplanted patients. Front. Microbiol.

Beaume, M., Köhler, T., Greub, G., Manuel, O., Aubert, J.D., Baerlocher, L., Farinelli, L., Buckling, A., and Van Delden, C. (2017). Rapid adaptation drives invasion of airway donor microbiota by Pseudomonas after lung transplantation. Sci. Rep.

Bernasconi, E., Pattaroni, C., Koutsokera, A., Pison, C., Kessler, R., Benden, C., Soccal, P.M., Magnan, A., Aubert, J.-D., Marsland, B.J., et al. (2016). Airway Microbiota Determines Innate Cell Inflammatory or Tissue Remodeling Profiles in Lung Transplantation. Am. J. Respir. Crit. Care Med. 194, 1252–1263.

Blatter, J.A., Sweet, S.C., Conrad, C., Danziger-Isakov, L.A., Faro, A., Goldfarb, S.B., Hayes, D., Melicoff, E., Schecter, M., Storch, G., et al. (2018). Anellovirus loads are associated with outcomes in pediatric lung transplantation. Pediatr. Transplant.

Borewicz, K., Pragman, A.A., Kim, H.B., Hertz, M., Wendt, C., and Isaacson, R.E. (2013). Longitudinal analysis of the lung microbiome in lung transplantation. FEMS Microbiol. Lett.

Bowers, E.F., and Jeffries, L.R. (1955). Optochin in the identification of str. pneumoniae. J. Clin. Pathol.

Brennan, A.L., Gyi, K.M., Wood, D.M., Johnson, J., Holliman, R., Baines, D.L., Philips, B.J., Geddes, D.M., Hodson, M.E., and Baker, E.H. (2007). Airway glucose concentrations and effect on growth of respiratory pathogens in cystic fibrosis. J. Cyst. Fibros.

Broggi, A., Granucci, F., and Zanoni, I. (2020). Type III interferons: Balancing tissue tolerance and resistance to pathogen invasion. J. Exp. Med. 217.

Caporaso, J.G., Kuczynski, J., Stombaugh, J., Bittinger, K., Bushman, F.D., Costello, E.K., Fierer, N., Pẽa, A.G., Goodrich, J.K., Gordon, J.I., et al. (2010). QIIME allows analysis of high-throughput community sequencing data. Nat. Methods.

Carney, S.M., Clemente, J.C., Cox, M.J., Dickson, R.P., Huang, Y.J., Kitsios, G.D., Kloepfer, K.M., Leung, J.M., LeVan, T.D., Molyneaux, P.L., et al. (2020). Methods in Lung Microbiome Research. Am. J. Respir. Cell Mol. Biol. 62, 283–299.

Charlson, E.S., Bittinger, K., Haas, A.R., Fitzgerald, A.S., Frank, I., Yadav, A., Bushman, F.D., and Collman, R.G. (2011). Topographical continuity of bacterial populations in the healthy human respiratory tract. Am. J. Respir. Crit. Care Med.

Charlson, E.S., Diamond, J.M., Bittinger, K., Fitzgerald, A.S., Yadav, A., Haas, A.R., Bushman, F.D., and Collman, R.G. (2012). Lung-enriched organisms and aberrant bacterial and fungal respiratory microbiota after lung transplant. Am. J. Respir. Crit. Care Med.

Chong, A.S., and Alegre, M.-L. (2014). Transplantation tolerance and its outcome during infections and inflammation. Immunol. Rev. 258, 80–101.

Cohen, T.S., Hilliard, J.J., Jones-Nelson, O., Keller, A.E., O’Day, T., Tkaczyk, C., Digiandomenico, A., Hamilton, M., Pelletier, M., Wang, Q., et al. (2016). Staphylococcus aureus α toxin potentiates opportunistic bacterial lung infections. Sci. Transl. Med.

Cummings, L.A., Hoogestraat, D.R., Rassoulian-Barrett, S.L., Rosenthal, C.A., Salipante, S.J., Cookson, B.T., and Hoffman, N.G. (2020). Comprehensive evaluation of complex polymicrobial specimens using next generation sequencing and standard microbiological culture. Sci. Rep. 10, 5446.

Dickson, R.P., and Huffnagle, G.B. (2015). The Lung Microbiome: New Principles for Respiratory Bacteriology in Health and Disease. PLOS Pathog. 11, e1004923.

Dickson, R.P., Erb-Downward, J.R., Freeman, C.M., McCloskey, L., Beck, J.M., Huffnagle, G.B., and Curtis, J.L. (2015). Spatial variation in the healthy human lung microbiome and the adapted island model of lung biogeography. Ann. Am. Thorac. Soc.

Dickson, R.P., Erb-Downward, J.R., Freeman, C.M., McCloskey, L., Falkowski, N.R., Huffnagle, G.B., and Curtis, J.L. (2017). Bacterial topography of the healthy human lower respiratory tract. MBio 8.

Erb-Downward, J.R., Thompson, D.L., Han, M.K., Freeman, C.M., McCloskey, L., Schmidt, L.A., Young, V.B., Toews, G.B., Curtis, J.L., Sundaram, B., et al. (2011). Analysis of the lung microbiome in the “healthy” smoker and in COPD. PLoS One.

Evans, C.R., Karnovsky, A., Kovach, M.A., Standiford, T.J., Burant, C.F., and Stringer, K.A. (2014). Untargeted LC-MS metabolomics of bronchoalveolar lavage fluid differentiates acute respiratory distress syndrome from health. J. Proteome Res. 13, 640–649.

Faure, K., Sawa, T., Ajayi, T., Fujimoto, J., Moriyama, K., Shime, N., and Wiener-Kronish, J.P. (2004). TLR4 signaling is essential for survival in acute lung injury induced by virulent Pseudomas aeruginosa secreting type III secretory toxins. Respir. Res.

Frye, L., and Machuzak, M. (2017). Airway Complications After Lung Transplantation. Clin. Chest Med. 38, 693–706.

Fusillo, M.H., and Weiss, D.L. (1959). Qualitative estimation of staphylococcal deoxyribonuclease. J. Bacteriol.

Geijtenbeek, T.B.H., and Gringhuis, S.I. (2009). Signalling through C-type lectin receptors: Shaping immune responses. Nat. Rev. Immunol.

Gill, S.K., Hui, K., Farne, H., Garnett, J.P., Baines, D.L., Moore, L.S.P., Holmes, A.H., Filloux, A., and Tregoning, J.S. (2016). Increased airway glucose increases airway bacterial load in hyperglycaemia. Sci. Rep. 6, 27636.

Görzer, I., Jaksch, P., Strassl, R., Klepetko, W., and Puchhammer-Stöckl, E. (2017). Association between plasma Torque teno virus level and chronic lung allograft dysfunction after lung transplantation. J. Hear. Lung Transplant. 36, 366–368.

Gregson, A.L., Wang, X., Weigt, S.S., Palchevskiy, V., Lynch, J.P., Ross, D.J., Kubak, B.M., Saggar, R., Fishbein, M.C., Ardehali, A., et al. (2013). Interaction between pseudomonas and CXC chemokines increases risk of bronchiolitis obliterans syndrome and death in lung transplantation. Am. J. Respir. Crit. Care Med.

Hardison, M.T., Galin, F.S., Calderon, C.E., Djekic, U. V., Parker, S.B., Wille, K.M., Jackson, P.L., Oster, R.A., Young, K.R., Blalock, J.E., et al. (2009). The Presence of a Matrix-Derived Neutrophil Chemoattractant in Bronchiolitis Obliterans Syndrome after Lung Transplantation. J. Immunol. 182, 4423–4431.

Hilty, M., Burke, C., Pedro, H., Cardenas, P., Bush, A., Bossley, C., Davies, J., Ervine, A., Poulter, L., Pachter, L., et al. (2010). Disordered microbial communities in asthmatic airways. PLoS One.

Ishii, T., Wallace, A.M., Zhang, X., Gosselink, J., Abboud, R.T., English, J.C., Paré, P.D., and Sandford, A.J. (2006). Stability of housekeeping genes in alveolar macrophages from COPD patients. Eur. Respir. J.

Ivanov, I.I., Frutos, R. de L., Manel, N., Yoshinaga, K., Rifkin, D.B., Sartor, R.B., Finlay, B.B., and Littman, D.R. (2008). Specific Microbiota Direct the Differentiation of IL-17-Producing T-Helper Cells in the Mucosa of the Small Intestine. Cell Host Microbe.

Jaksch, P., Kundi, M., Görzer, I., Muraközy, G., Lambers, C., Benazzo, A., Hoetzenecker, K., Klepetko, W., and Puchhammer-Stöckl, E. (2018). Torque Teno Virus as a Novel Biomarker Targeting the Efficacy of Immunosuppression After Lung Transplantation. J. Infect. Dis. 218, 1922–1928.

Jorth, P., Staudinger, B.J., Wu, X., Hisert, K.B., Hayden, H., Garudathri, J., Harding, C.L., Radey, M.C., Rezayat, A., Bautista, G., et al. (2015). Regional Isolation Drives Bacterial Diversification within Cystic Fibrosis Lungs. Cell Host Microbe.

Kelly, F.J., and Mudway, I.S. (2003). Protein oxidation at the air-lung interface. Amino Acids 25, 375–396.

Kešnerová, L., Emery, O., Troilo, M., Liberti, J., Erkosar, B., and Engel, P. (2020). Gut microbiota structure differs between honeybees in winter and summer. ISME J. 14, 801–814.

Koutsokera, A., Royer, P.J., Antonietti, J.P., Fritz, A., Benden, C., Aubert, J.D., Tissot, A., Botturi, K., Roux, A., Reynaud-Gaubert, M.L., et al. (2017). Development of a multivariate prediction model for early-onset bronchiolitis obliterans syndrome and restrictive allograft syndrome in lung transplantation. Front. Med.

Kursa, M.B., and Rudnicki, W.R. (2010). Feature selection with the boruta package. J. Stat. Softw.

Lloyd, C.M., and Marsland, B.J. (2017). Lung Homeostasis: Influence of Age, Microbes, and the Immune System. Immunity 46, 549–561.

Mallia, P., Webber, J., Gill, S.K., Trujillo-Torralbo, M.-B., Calderazzo, M.A., Finney, L., Bakhsoliani, E., Farne, H., Singanayagam, A., Footitt, J., et al. (2018). Role of airway glucose in bacterial infections in patients with chronic obstructive pulmonary disease. J. Allergy Clin. Immunol. 142, 815–823.e6.

Marsland, B.J., and Gollwitzer, E.S. (2014). Host–microorganism interactions in lung diseases. Nat. Rev. Immunol. 14, 827–835.

Martinu, T., Pavlisko, E.N., Chen, D.F., and Palmer, S.M. (2011). Acute Allograft Rejection: Cellular and Humoral Processes. Clin. Chest Med.

McMurdie, P.J., and Holmes, S. (2013). Phyloseq: An R Package for Reproducible Interactive Analysis and Graphics of Microbiome Census Data. PLoS One.

Mika, M., Nita, I., Morf, L., Qi, W., Beyeler, S., Bernasconi, E., Marsland, B.J., Ott, S.R., von Garnier, C., and Hilty, M. (2018). Microbial and host immune factors as drivers of COPD. ERJ Open Res. 4, 00015–02018.

Molyneaux, P.L., Cox, M.J., Willis-Owen, S.A.G., Mallia, P., Russell, K.E., Russell, A.M., Murphy, E., Johnston, S.L., Schwartz, D.A., Wells, A.U., et al. (2014). The role of bacteria in the pathogenesis and progression of idiopathic pulmonary fibrosis. Am. J. Respir. Crit. Care Med.

Mouraux, S., Bernasconi, E., Pattaroni, C., Koutsokera, A., Aubert, J.D., Claustre, J., Pison, C., Royer, P.J., Magnan, A., Kessler, R., et al. (2017). Airway microbiota signals anabolic and catabolic remodeling in the transplanted lung. J. Allergy Clin. Immunol.

Ng, T.H.S., Britton, G.J., Hill, E. V., Verhagen, J., Burton, B.R., and Wraith, D.C. (2013). Regulation of adaptive immunity; the role of interleukin-10. Front. Immunol.

Nosotti, M., Tarsia, P., and Morlacchi, L.C. (2018). Infections after lung transplantation. J. Thorac. Dis.

Odendall, C., Voak, A.A., and Kagan, J.C. (2017). Type III IFNs Are Commonly Induced by Bacteria-Sensing TLRs and Reinforce Epithelial Barriers during Infection. J. Immunol. 199, 3270–3279.

Pattaroni, C., Watzenboeck, M.L., Schneidegger, S., Kieser, S., Wong, N.C., Bernasconi, E., Pernot, J., Mercier, L., Knapp, S., Nicod, L.P., et al. (2018). Early-Life Formation of the Microbial and Immunological Environment of the Human Airways. Cell Host Microbe 24, 857–865.e4.

Reynolds, A.P., Richards, G., de la Iglesia, B., and Rayward-Smith, V.J. (2006). Clustering Rules: A Comparison of Partitioning and Hierarchical Clustering Algorithms. J. Math. Model. Algorithms 5, 475–504.

Rognes, T., Flouri, T., Nichols, B., Quince, C., and Mahé, F. (2016). VSEARCH: a versatile open source tool for metagenomics. PeerJ.

Schubert, E., and Rousseeuw, P.J. (2019). Faster k-Medoids Clustering: Improving the PAM, CLARA, and CLARANS Algorithms. In Lecture Notes in Computer Science (Including Subseries Lecture Notes in Artificial Intelligence and Lecture Notes in Bioinformatics), p.

Segal, L.N., Alekseyenko, A. V., Clemente, J.C., Kulkarni, R., Wu, B., Chen, H., Berger, K.I., Goldring, R.M., Rom, W.N., Blaser, M.J., et al. (2013). Enrichment of lung microbiome with supraglottic taxa is associated with increased pulmonary inflammation. Microbiome 1, 19.

Segal, L.N., Clemente, J.C., Tsay, J.C.J., Koralov, S.B., Keller, B.C., Wu, B.G., Li, Y., Shen, N., Ghedin, E., Morris, A., et al. (2016). Enrichment of the lung microbiome with oral taxa is associated with lung inflammation of a Th17 phenotype. Nat. Microbiol.

Segura-Wang, M., Görzer, I., Jaksch, P., and Puchhammer-Stöckl, E. (2018). Temporal dynamics of the lung and plasma viromes in lung transplant recipients. PLoS One 13, e0200428.

Shimazu, R., Akashi, S., Ogata, H., Nagai, Y., Fukudome, K., Miyake, K., and Kimoto, M. (1999). MD-2, a molecule that confers lipopolysaccharide responsiveness on toll-like receptor 4. J. Exp. Med.

Simon-Soro, A., Sohn, M.B., Mcginniss, J.E., Imai, I., Brown, M.C., Knecht, V.R., Bailey, A., Clarke, E.L., Cantu, E., Li, H., et al. (2019). Upper respiratory dysbiosis with a facultative-dominated ecotype in advanced lung disease and dynamic change after lung transplant. Ann. Am. Thorac. Soc.

Simon, A.R., Takahashi, S., Severgnini, M., Fanburg, B.L., and Cochran, B.H. (2002). Role of the JAK-STAT pathway in PDGF-stimulated proliferation of human airway smooth muscle cells. Am. J. Physiol. - Lung Cell. Mol. Physiol.

Todd, J.L., Kelly, F.L., Nagler, A., Banner, K., Pavlisko, E.N., Belperio, J.A., Brass, D., Weigt, S.S., and Palmer, S.M. (2020). Amphiregulin contributes to airway remodeling in chronic allograft dysfunction after lung transplantation. Am. J. Transplant. 20, 825–833.

Vandeputte, D., Kathagen, G., D’hoe, K., Vieira-Silva, S., Valles-Colomer, M., Sabino, J., Wang, J., Tito, R.Y., De Commer, L., Darzi, Y., et al. (2017). Quantitative microbiome profiling links gut community variation to microbial load. Nature 551, 507–511.

Venkataraman, A., Bassis, C.M., Beck, J.M., Young, V.B., Curtis, J.L., Huffnagle, G.B., and Schmidt, T.M. (2015). Application of a neutral community model to assess structuring of the human lung microbiome. MBio.

Verleden, S.E., Von, J., der Thüsen, Roux, A., Brouwers, E.S., Braubach, P., Kuehnel, M., Laenger, F., and Jonigk, D. (2020). When tissue is the issue: a histological review of chronic lung allograft dysfunction. Am. J. Transplant. ajt.15864.

De Vlaminck, I., Khush, K.K., Strehl, C., Kohli, B., Luikart, H., Neff, N.F., Okamoto, J., Snyder, T.M., Cornfield, D.N., Nicolls, M.R., et al. (2013). Temporal response of the human virome to immunosuppression and antiviral therapy. Cell.

Whelan, F.J., Waddell, B., Syed, S.A., Shekarriz, S., Rabin, H.R., Parkins, M.D., and Surette, M.G. (2020). Culture-enriched metagenomic sequencing enables in-depth profiling of the cystic fibrosis lung microbiota. Nat. Microbiol. 5, 379–390.

Willner, D.L., Hugenholtz, P., Yerkovich, S.T., Tan, M.E., Daly, J.N., Lachner, N., Hopkins, P.M., and Chambers, D.C. (2013). Reestablishment of recipient-associated microbiota in the lung allograft is linked to reduced risk of bronchiolitis obliterans syndrome. Am. J. Respir. Crit. Care Med.

Winstanley, C., O’Brien, S., and Brockhurst, M.A. (2016). Pseudomonas aeruginosa Evolutionary Adaptation and Diversification in Cystic Fibrosis Chronic Lung Infections. Trends Microbiol.

Young, J.C., Chehoud, C., Bittinger, K., Bailey, A., Diamond, J.M., Cantu, E., Haas, A.R., Abbas, A., Frye, L., Christie, J.D., et al. (2015). Viral Metagenomics Reveal Blooms of Anelloviruses in the Respiratory Tract of Lung Transplant Recipients. Am. J. Transplant. 15, 200–209.

Zaneveld, J.R., McMinds, R., and Thurber, R.V. (2017). Stress and stability: Applying the Anna Karenina principle to animal microbiomes. Nat. Microbiol.

